# Neural dynamics underlying minute-timescale persistent behavior in the human brain

**DOI:** 10.1101/2024.07.16.603717

**Authors:** Hristos S. Courellis, Taufik A. Valiante, Adam N. Mamelak, Ralph Adolphs, Ueli Rutishauser

## Abstract

The ability to pursue long-term goals relies on a representations of task context that can both be maintained over long periods of time and switched flexibly when goals change. Little is known about the neural substrate for such minute-scale maintenance of task sets. Utilizing recordings in neurosurgical patients, we examined how groups of neurons in the human medial frontal cortex and hippocampus represent task contexts. When cued explicitly, task context was encoded in both brain areas and changed rapidly at task boundaries. Hippocampus exhibited a temporally dynamic code with fast decorrelation over time, preventing cross-temporal generalization. Medial frontal cortex exhibited a static code that decorrelated slowly, allowing generalization across minutes of time. When task context needed to be inferred as a latent variable, hippocampus encoded task context with a static code. These findings reveal two possible regimes for encoding minute-scale task-context representations that were engaged differently based on task demands.

## Introduction

We can engage in persistent behavior for extended periods of time that span from seconds to years. For example, a single question conveyed over the course of a few seconds could lead to a 2-minute trip to the kitchen or a 20-minute quest to find lost keys. Such temporally extended goal-directed behavior is thought to rely on persistent representations of task context^1–5^ (also referred to as task sets). Task contexts specify the structure and goal of the task being pursued and include rules specifying which actions to take, the meaning of different stimuli, instructed or learned associations between stimuli, actions, and rewards. In humans, task context is typically specified explicitly through symbolic or language-based instructions^6,7^. Alternatively, task context can also be inferred implicitly from the temporal statistics of the environment ^8–10^ and/or trial-and error learning^11–14^. Most tasks are learned this way by animals. A major open question is how task context is maintained in the brain over time periods of minutes to hours, and how such representations enable persistent behavior.

During working memory maintenance, subsets of cells in the human medial temporal lobe (MTL) remain persistently active if their preferred stimulus is held in short-term memory^15,16–18^. Similarly, in monkeys and rodents, cells in the frontal and parietal lobe have been described that signal remembered stimulus identities, spatial locations, and/or motor plans during the delay periods of working memory tasks^19–21^. The activity of these cells predicts working memory content and quality, suggesting that they are part of the neural substrate of working memory. A common feature of working memory tasks is that stimuli are maintained for a brief period of time (typically 1-3 seconds). Here, in contrast, we examine how task context is maintained over periods of 1-10 minutes, which is an order of magnitude longer. Little is known about the neural substrate for working memories maintained over such extended periods of time.

A second open question relates to the format in which long-time period task context information is represented. During the brief maintenance time periods of working memory task, two types of representations have been found: static and dynamic. Static representations are such that the same neurons represent the same information throughout the maintenance period. In dynamic representations, on the other hand, a given neuron’s tuning changes throughout the maintenance period. Static representations allow generalizing behavior flexibly across arbitrarily long periods of time^22,23^ and to new stimuli^24^ and would therefore seem well suited for representing task context. Dynamic representations, on the other hand, have advantages in terms of the encoding of time and the avoidance of interference between sensory inputs and information held in memory (among others) ^25–27^. Here, we examine whether representations of task context are flexible or dynamic using cross-temporal generalization analysis.

Our focus in this study is on two areas of the human brain that have been implicated in the maintenance of task context: the medial frontal cortex (MFC) and the hippocampus. In the MFC, we are examining neural responses in the dorsal Anterior Cingulate Cortex (dACC) and the pre-Supplementary Motor Area (preSMA), which are thought to be critical for flexible temporally extended behavior^28^. MFC represents task sets^29^, guides the task-dependent selection of appropriate actions, and mediates the switching between action sets^30^. Indeed, MFC is necessary for successful persistent behavior given the effects of lesions in this area of the brain^31,32^. The hippocampus, on the other hand, is thought to combine contextual variables with information about time and sensory stimuli to form cognitive maps and to situate sensory inputs contextually. This view is, for example, supported by the presence of time cells and ramp cells, whose activity encodes the passage of time ^33–36^ so as to give rise to a ‘temporal context’ that enables unique episodic memories^37^. How hippocampal neurons represent a continually changing temporal context while simultaneously also representing temporally static task context remains an open question. Given this prior literature, we hypothesize that (i) the MFC harbors long-time scale persistent representations of task context, and that (ii) the hippocampus encodes context more transiently in a dynamic fashion.

## Results

To examine these hypotheses, we utilized human single neuron recordings from epilepsy patients who performed two cognitive experiments (Experiment 1 and 2, see methods) that required the maintenance of task instructions over long periods of time ^6,12^. The work described here is new analysis of these two previously published data sets^6^.

In Experiment 1, n=13 patients completed 33 sessions of a blocked context-dependent decision-making task. Patients were shown a sequence of images and were asked to make binary “Yes vs. No” decisions for each image shown (Fig. 1A). The question (task context) to be answered in a given block was either a semantic categorization question (“Is the image a member of category X?”) or a recognition memory question (“Have you seen this image before?”). Task contexts were shown on the screen only once at the beginning of every block and needed to be remembered for the following 40 trials in the block (mean block duration 115.85s ± 4.31s, mean ± s.e.m.). Thus, successfully performing this task required memory for the task context for up to 2 minutes. Patients completed 320 trials (8 blocks) in each session (total duration 1100.0s ± 37.1s, mean ± s.e.m.). The first block was always a categorization block and every ensuing block alternated between memory and categorization (Fig. 1A, bottom). Each trial consisted of a pre-stimulus baseline, followed by presentation of the stimulus, which was displayed until patients provided a response (Fig. 1A, top). No trial-by-trial feedback was provided. Patients performed the task with high accuracy (83.6% ± 1.1%, mean ± s.e.m. over n=33 sessions) and speed (reaction time relative to stimulus onset was 1.34s ± 0.10s, mean ± s.e.m. over trials).

**Figure 1.**
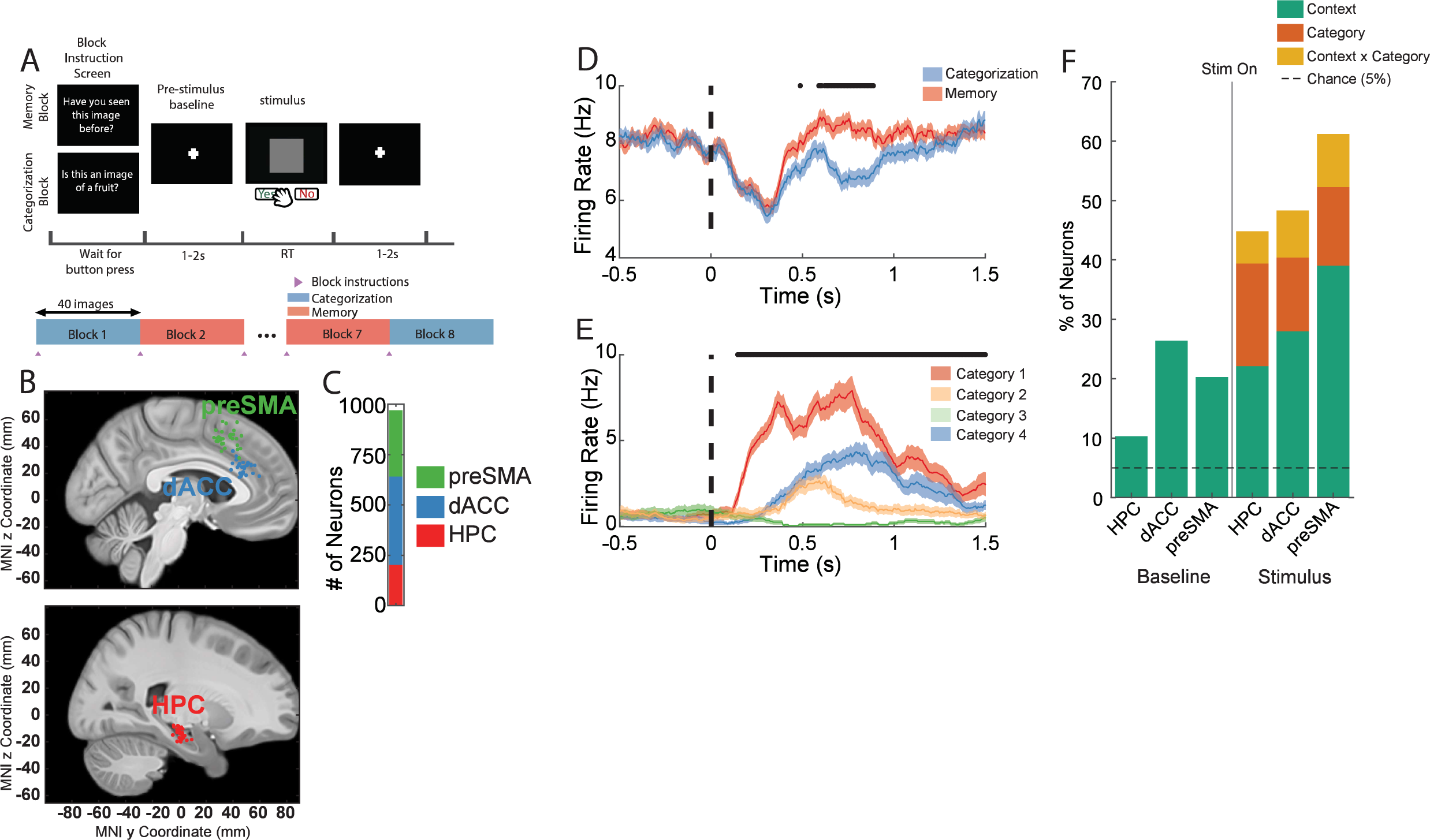
Single neurons are tuned to task variables during instructed task switching. **(A)** Experiment 1 consisted of eight blocks of 40 trials each. Task context alternated between a categorization task and a recognition memory task. Instructions were provided to patients only once at the start of each block but applied for all trials until the next set of instructions. **(B)** Electrode locations for the pre-Supplementary Motor Area (preSMA, green), dorsal Anterior Cingulate Cortex (dACC, blue), and anterior hippocampus (HPC, red). Each dot corresponds to the implant site of a microwire bundle for a single patient. All implants were bilateral, and electrodes are shown on the same hemisphere for visualization purposes. **(C)** Number of single units recorded across the three brain areas (970 neurons total, dACC, blue = 329, preSMA, green = 438, HPC, red = 203). **(D-E)** Example PSTHs of neurons recorded in HPC that are differentially selective for task context **(D)** and semantic category of the visual stimulus **(E)** during stimulus presentation throughout the task. Stimulus onset occurs at time 0. Black points above the PSTH indicate times where a sliding-window 1-way ANOVA (250 msec width) over the considered task variables was significant (p < 0.05).**(F)** Percentage of neurons that exhibit tuning to task variables during the Baseline (-1 to 0s prior to stimulus onset) and Stimulus periods (0.2 to 1.2 following stimulus onset). Neurons are considered tuned during the stimulus period to either a main effect (context – green, category – orange), or the interaction (orange) if the associated factor in a 2x4 ANOVA (Context x Category) was significant (p<0.05). Baseline tuning is determined using a 1-way ANOVA for context. Horizontal dashed line indicates chance level.

### The hippocampal context representation is temporally dynamic, whereas MFC is static

Across all sessions, we recorded in total n=970 single neurons (Fig. 1B,C) from the Hippocampus (HPC, 203 neurons), dorsal Anterior Cingulate Cortex (dACC, 329 neurons), and pre-Supplementary Motor Area (preSMA, 438 neurons). At the single neuron level, the response of significant proportions of neurons was modulated by task context both during the baseline (-1s to 0s prior to stimulus onset) and the stimulus period (0.2s to 1.2s after stimulus onset) in all three brain areas (Fig. 1D shows an example neuron and Fig. 1F shows statistics; p < 0.05, 1-Way ANOVA for context during baseline). In addition, following stimulus onset, neurons were modulated by semantic image category alone or interactions between category and task context in all three brain areas (Fig. 1E shows an example neuron and Fig. 1F shows statistics; 2-Way ANOVA for context and category during stimulus). Additional example neurons encoding task context during the baseline and stimulus periods from all three regions are shown in Fig. S1. Thus, at the single-neuron level, there are neurons whose firing rate is significantly modulated by the task context on average across the entire experiment.

We started examining the format of the task context representations formed by conducting population-level analysis. We constructed pseudopopulations of neurons in each region pooled across sessions, and trained linear support vector machines (SVMs) to decode task context on individual trials during the stimulus and baseline periods (Fig. 2A, inset, see Methods for details). Across all trials, task context was significantly decodable in all three brain areas during both the stimulus and baseline periods: HPC (Chance is 50%. 60.3% *base*, *p*_*base*_ = 0.007, 71.5% *stim*, *p*_*stim*_ = 9.5×10^−6^), dACC (83.9% *base*, *p*_*base*_ = 1.4×10^−13^, 91.7% *stim*, *p*_*stim*_ = 0), and preSMA(82.9% *base*, *p*_*base*_ = 6.1×10^−15^, 99.9% *stim*, *p*_*stim*_ = 0). All reported p-values are computed using a non-parametric permutation test based on a null distribution estimated by retraining decoders after trial-label shuffling. Only correct trials (subject provided correct answer) were used for decoder training/testing, and all decoders were matched for number of neurons and number of trials per condition through random sub-sampling unless otherwise specified.

**Figure 2.**
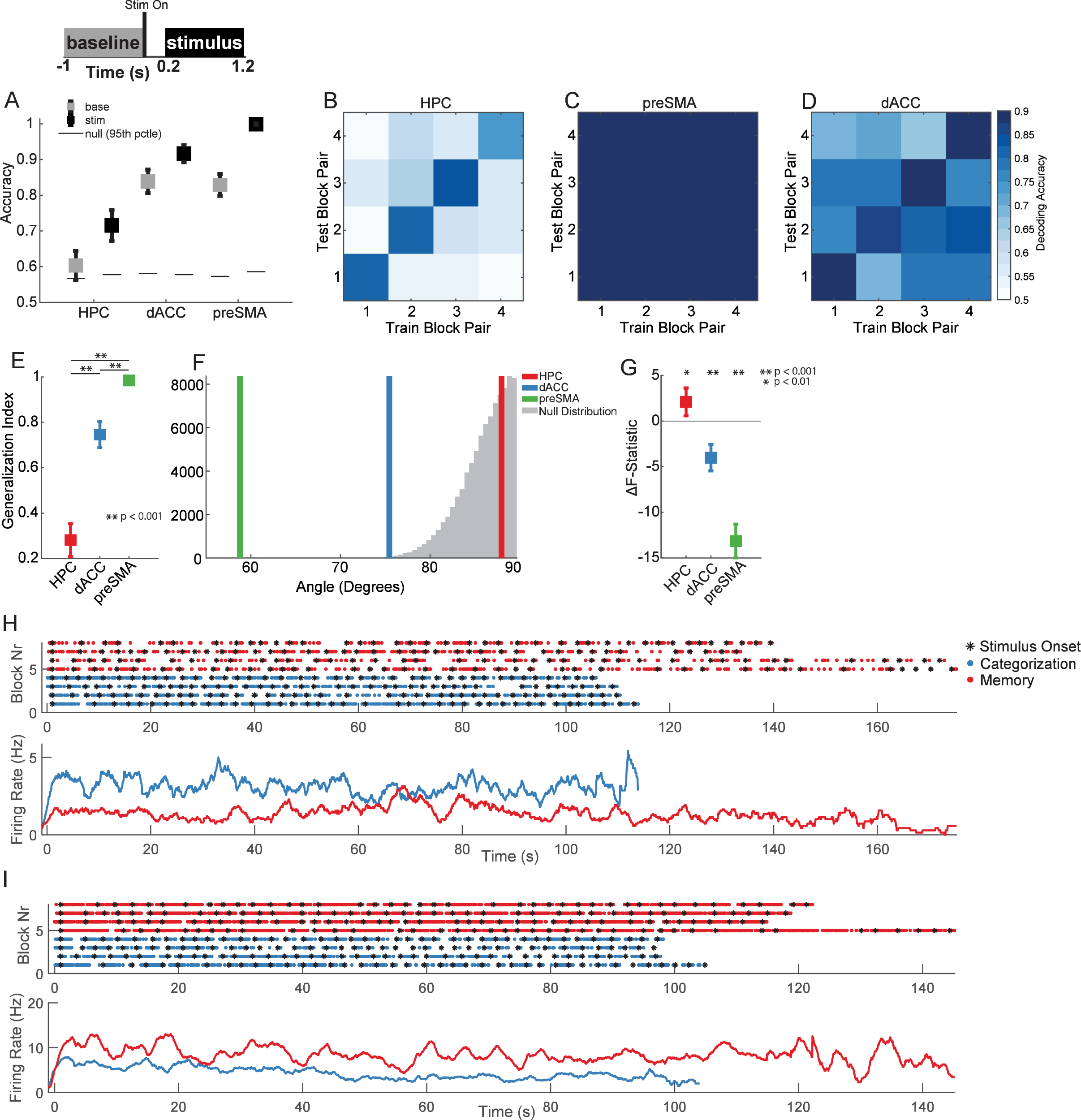
Task context representation temporally generalizes in MFC but not HPC. **(A)** Context decoding accuracy during the baseline (gray) and stimulus (black) periods (see inset) using correct trials from the entire experiment. Black horizontal lines indicate 95^th^ percentile of null distribution. Chance is 0.5. **(B-D)** Cross-temporal decoding. Plots indicating the out-of-sample decoding accuracy of decoders trained to decode context from correct trials in adjacent block pairs during the stimulus period. On-diagonal decoding accuracies (train/test on same block pair) are reported with 5-fold cross validation. Off-diagonal decoding accuracies use all available trials for training and testing. The colormap shown applies to all three panels. **(E)** Generalization index, quantifying the cross-temporal generalization of context decoding across block pairs. Index values range from 0 to 1, indicating no generalization and perfect generalization of context coding respectively. P-values are computed using two-tailed Wilcoxon Rank-sum tests. **(F)** Angles computed between vectors normal to the hyperplanes of the block-pair context decoders. Angles here were estimated in an n = 150-dimensional space to facilitate direct comparison of angle values between regions. The mean angle averaged over all pairs of decoders is reported for HPC (red), dACC (blue), and preSMA (green). The null distribution (gray) is populated by the angle between randomly selected pairs of trial-shuffled context decoders. **(G)** Comparison of ANOVA F-statistics fit using block number and task context in single-neurons. Values are reported are mean ± s.e.m. ΔF-statistic computed over neurons. P-values are computed using a two-sided t-test. **(H)** Example raster (above) and PSTH (below) for a neuron in the dACC that exhibited persistent firing rate context modulation throughout entire blocks. An individual row in the raster (above) corresponds to the activity of a single neuron plotted for a block. Each point corresponds to one spike discharged by the neuron. Black stars indicate stimulus onset times. Blocks are re-ordered according to task context (categorization = blue, memory = red), and are aligned to the stimulus onset time of the first trial in each block. PSTH (below) shows mean firing rate computed over blocks. **(I)** Same as **(H)**, but for a neuron in preSMA. In all panels, plots with squares and error bars indicate mean ± s.e.m. of the computed metric (e.g. decoding accuracy, generalization index, etc…) over 250 iterations of bootstrapped re-sampling unless otherwise specified (see methods). Null distributions were computed with 250 iterations of trial-label shuffling followed by re-computing the metric in question.

We next assessed how stable the representation of task context was throughout each session using cross-temporal generalization analysis. If the representation were static, decoders trained to discriminate task context during one part of the experimental session should generalize to other parts of the session. If, on the other hand, the task context representation were dynamic, then context decoders trained in one part of the session should perform poorly when generalized to other parts of the session. Context decoders were trained on adjacent block pairs so that one block of each kind is present in each pair. We used the neural responses following stimulus onset for this analysis (since context representations are strongest during this time period in all brain areas, see Fig. 2A). In the hippocampus, task context decoding did not generalize across time during stimulus processing in HPC (Fig. 2B; as indicated by higher on-diagonal vs. off-diagonal decoding accuracy). In contrast, task context decoding generalized across time in both the parts of the MFC (Fig. 2C-D), with high decoding accuracy both on and off-diagonal (see below for quantification). This impression was confirmed quantitatively: the generalization index for both dACC (Fig. 2E, blue vs red, *p* < 0.001, Permutation test) and preSMA (Fig. 2E, blue vs red, *p* < 0.001, Permutation test) was close to 1 (1 is maximal generalization and 0 is no generalization) and significantly larger than in the HPC.

To distinguish between instances where the decoders relied on few and many task-context modulated neurons to generalize, we also computed the angle between context coding vectors in each block pair. We define the coding vector here to be the vector of importance indices assigned to each neuron during training. Whereas for decoding accuracy, a decoder could achieve equally high performance by relying on a small number of well-tuned features or a large number of more weakly tuned features, the angle between coding vectors can only differ from orthogonal in a high dimensional space (n=150 dimensions here) if many neurons are contributing to decoder performance. Pairs of context coding vectors between any two block pairs significantly differ from orthogonal both during the stimulus period for dACC (Fig. 2F, blue, 76.5°, *p* < 0.001 against shuffle null) and preSMA (Fig. 2F, green, 59.0°, *p* < 0.001 against shuffle null), indicating significant context coding vector alignment across block pairs. In HPC, on the other hand, context coding vector angles did not significantly differ from orthogonal across block pairs (Fig. 2F, red, 88.5°, *p* = 0.46 against shuffle null). Note that angles here are computed in a 150-dimensional space constructed by sub-sampling neurons randomly over iterations. All of the above cross-temporal context generalization findings were also present during the baseline period for the three regions (Fig. S2), with the notable exception that the cross-block pair generalization index did not significantly differ between dACC and preSMA (Fig. S2D, blue vs green).

Together, these findings indicate that the code for task context is dynamic in the HPC. In contrast, in the MFC, context coding is static. What about the neuronal responses in the HPC causes the code to be dynamic? One possibility is that individual HPC neurons do not reliably represent a given task context throughout a session. This stands in contrast to preSMA and dACC, where generalizing context decoders imply that single neurons in these regions represent task context with a static rate code. To test this prediction, we fit 2-Way ANOVAs on all recorded neurons in a given brain area individually with two categorical regressors for block number and task context. We reasoned that for activity of neurons supporting cross-temporal generalization, the main effect of task context would explain more variance than the block number main effect (and vice versa for neurons supporting a dynamic code). We therefore next compared the amount of variance explained by the two main effects (ΔF-statistic, see methods). Variance in single-neuron responses in the hippocampus was on average better explained by the block-specific regressor during both stimulus (Fig 2G, red, *p* = 0.017, Student’s t-test) and baseline (Fig S2F, red, *p* = 0.02, Student’s t-test) periods. In dACC and preSMA, on the other hand, the task-context regressor explained significantly more variance during the stimulus (Fig 2G, *p*_*green*_, *p*_*blue*_ < 0.001), but not during the baseline (Fig S2F, *p*_*green*_, *p*_*blue*_ > 0.01) period, though the general trend remained. Thus, the cross-block pair generalization pattern in these regions can be accounted for by different encoding strategies at the level of single-units. These differences in encoding strategy can be appreciated by viewing the activity of a given neuron across the entire block, revealing that dACC (e.g. Fig. 2H, S3A) and preSMA (e.g. Fig. 2I,S3B,C) neurons exhibit task context modulated firing that persists for the duration of entire blocks. In contrast, no such neurons are present in HPC (e.g. Fig. S3D-F).

### Relationship with image category representations and temporal stability

The dynamic code for task context on the timescale of blocks in the hippocampus raises several questions. First, were shifts in the context representation over time driven by abrupt changes at the beginning of each block, leaving a static context representation within each block? To address this question, we performed a block-half decoding and generalization analysis where decoders for task context were trained using data from the first half and second half of every block, then evaluated on the second half and first half respectively. If the hippocampal context representation exhibited within-block dynamics, then first-and second-block half decoders should fail to generalize. If, however, the context-code remains static within individual blocks, then a context decoder trained on the first block half should do approximately equally well on the second block half (and vice versa from second to first). We find that for all brain areas, context decoders trained on one block half generalized well to the other block half during both the stimulus period (Fig. S4A-C) and the baseline period (Fig. S4D-F), with generalization indices close to 1 (Fig.S4C,F). This data suggests that the dynamic code in the HPC is due to changes that occur at the transition between blocks rather within blocks.

Second, can the temporal stability/instability of the context representation in different regions be explained by tuning strength differences of single-neurons and/or by recording instability? To address this possibility, we matched the distribution of single neuron ANOVA F-statistics for the main effect of context across the three regions before re-computing cross-block pair context decoding (see Methods for details). Following distribution matching, 174 neurons remained in each area for the subsequent cross-block pair analysis. This analysis revealed that the cross-block pair generalization indices for context remain qualitatively unchanged for the three regions during both the stimulus (Fig. S4G,H) and baseline (Fig. S4I,J) periods. The stability of the context code in dACC and preSMA can thus not be explained by stronger univariate context coding.

To demonstrate that the temporal stability of the context code in each region was robust to variation in trial and block duration within the experiment, we also analyzed a second variant of this experiment (6 sessions, 36 HPC, 75 dACC, 87 preSMA neurons). This variant of the experiment was longer due to requiring patients to wait for a 0.5-1.5-second period following stimulus offset till making a response. Mean trial duration was 2.47s ± 0.04s (vs 1.08s ± 0.05s in non-control session) and mean block duration was 162.54s ± 1.83s (vs 105.47s ± 2.23s in the original experiment). Despite the ∼250% increase in trial duration and ∼60% increase in block duration, dACC and preSMA exhibited significantly greater cross-block pair context stability both during stimulus (Fig. S4K) and baseline (Fig. S4L) periods compared to the HPC.

Lastly, to address recording stability, we analyze the geometry of a second task variable, image category. Image category is known from prior work to be encoded by HPC neurons in a static manner^7,16,17^ and can thus serve as a control for potential recording instability. Image category was decodable from all three regions, most prominently in the HPC (59.5%, *p* = 0, against shuffle null; chance=25%, Fig. S5A,B). Image category decoding generalized well across time in the HPC (Fig. S5C-G). Since in the same group of HPC neurons image category is encoded in a static manner, the temporally dynamic encoding of task context in the HPC is unlikely to be a result of recording instability.

### Hippocampal neural population exhibits faster temporal dynamics than MFC

The hippocampus contributes to episodic memory through the representation of temporal context^37–39^. One way to give rise to temporal context representations is through slow gradual drifts in neural population activity. We next asked whether such slowly changing temporal context representations were present in our data, and if so, whether they gave rise to our finding of dynamic encoding of task context. We examined neural population dynamics with single-trial resolution over the timescale of the experiment (∼20 mins) in a decoder-agnostic manner using the autocorrelation of the population response (see methods). This revealed a striking pattern: during the stimulus period, the HPC neural population response gradually and continually decorrelated as indicated by positive near-diagonal population vector correlations and increasingly negative correlations with increasing trial distance (Fig. 3A). In contrast, in the preSMA, a checkerboard-like autocorrelation pattern was present. This was due to trials with the same task context exhibiting positive correlations and trials in different task contexts exhibiting negative correlations across the entire experiment (Fig. 3C). A qualitatively similar pattern was present in the dACC (Fig. S6E). The same analysis performed during the baseline period revealed qualitatively similar, but less visually pronounced results in all three areas (Fig. S6A,C,G). These findings were also visible at the level of a single patient (Fig. S7).

**Figure 3.**
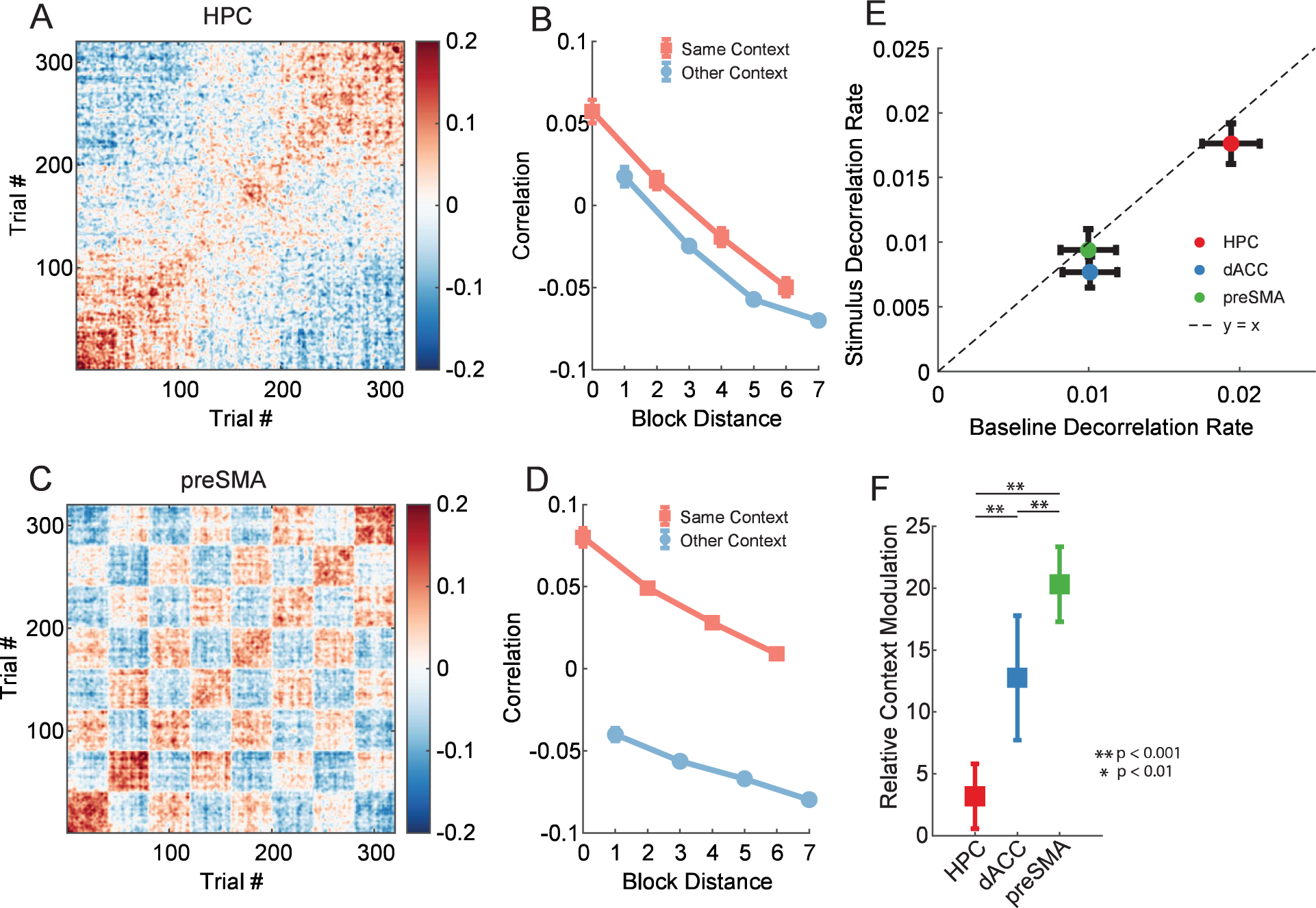
Hippocampal neural population exhibits temporal decorrelation. **(A)** Hippocampal population-vector autocorrelation matrix showing Pearson correlation for all possible pairs of trials. Correlations are computed between pseudo-population firing rate vectors computed for each trial. Diagonal values are removed for visualization purposes. Matrix shown here is an average over 250 iterations of sub-sampled estimation to match the number of neurons between regions. **(B)** Mean cross-block correlation computed over all possible pairs of trials that are in increasingly distant blocks for hippocampal neurons. For example, Block Distance 0 reports the average population-vector correlation between all pairs of trials in the same block, Block Distance 1 reports the average correlation between all pairs of trials exactly one block apart, etc. Even block distances correspond to blocks of the same task (light red), odd block distances correspond to blocks of the opposite task (light blue). Values are reported as mean ± s.e.m. over trial pairs. **(C,D)** Same as (**A,B)**, but for pre-SMA. **(E)** Baseline vs Stimulus population decorrelation rate plotted for each of the three regions. Decorrelation rate is estimated as the absolute value of the slope of the least-square fit to the cross-block decorrelation curves, e.g. shown in **(B)** and **(D)**. All slopes were negative, so increasing values indicate increasing rate of decorrelation with block distance. Circles and error bars correspond to mean and s.e.m. decorrelation rate computed over iterations of neuron sub-sampling. **(F)** Relative context modulation (cross-context correlation difference normalized by decorrelation rate) reported for the three areas during the stimulus period. Values are reported as mean ± s.e.m. over iterations of decorrelation curve estimation. P-values are computed by permutation test.

We next assessed how quickly the neural responses changed as a function of block distance by computing decorrelation curves (see methods). The population response became less correlated as a function of time in all brain areas (Fig. 3B,D) during both the stimulus and baseline period (see Fig. S6 for dACC). However, the HPC neural population decorrelated (see methods) significantly more rapidly than the dACC and preSMA during both the stimulus and baseline periods (Fig. 3E, red vs green and red vs blue, *p*_*stim*_ < 0.001, *p*_*base*_ < 0.001 pairwise two-tailed RankSum). Decorrelation speed did not differ significantly between dACC and preSMA (Fig. 3E, blue vs green, *p*_*stim*_ > 0.05, *p*_*base*_ > 0.05 RankSum). Also, decorrelation rate did not differ significantly between the categorization and memory task contexts for any region during both stimulus and baseline periods (Fig. S6I-N, blue vs red, all *p* > 0.05 RankSum).

Did switching between tasks cause changes in neural representations beyond those expected by the gradual decorrelation? If task switches had no effect, the task context representation of blocks of the same task (two blocks apart) would be twice as decorrelated as the representation of the opposite task (one block apart). To test whether this was the case, we computed the relative context modulation, which we defined as the average reduction in correlation from block distance 0 to block distance 1 normalized by the decorrelation rate.

Relative context modulation is a unitless indicator of the degree to which explicit changes in task context shift the neural representation while accounting for simultaneously occurring decorrelation (see Methods for details). We found that all three areas significantly differed from each other in their relative context modulation, with the preSMA and dACC exhibiting stronger context modulation effects (Fig. 3F, green, RCM = 20.0, blue, RCM = 12.8) and the HPC exhibiting the weakest effect (Fig. 3F, red, RCM = 3.2). These values indicate that the effect of task context switching is ∼3 times greater at driving changes in the HPC neural population as intrinsic decorrelation, whereas in medial frontal cortical structures the task-switching is ∼10-20 times stronger. A similar qualitative pattern was observed during the baseline, but with the dACC exhibiting greater relative context modulation than the preSMA (Fig. S6O). Thus, while the decorrelation rate of the hippocampal neural population is high, it exists alongside an even larger change in task context that is caused by the switching between tasks. Decorrelation alone can therefore not explain the cross-temporal context instability we find in our data in the HPC.

### The dACC context representation generalizes between stimulus and baseline periods

So far, we examined the dynamics of task context representations over time periods of blocks of 40 trials. We next turned to the shorter time period of a single trial, i.e. over about 2s. To do so, we asked what the relationship was between representations of task context during the baseline and stimulus periods within a given trial. At the single neuron level, some neurons were tuned to context in the same way during both time periods (Fig. 4A shows an example), others were tuned to context only in one of the time periods (e.g. Fig. S1A-D), and others were tuned but with inverse direction (e.g. Fig. S1E-F). The fraction of neurons tuned in both time periods (1-way ANOVA, p<0.05 separately in both time periods) was greater than would be expected by chance in all areas (Fig. 4B, purple, HPC = 18.0%, 11/61 neurons, dACC = 19.1%, 25/131 neurons, preSMA = 15.8%, 32/203 neurons, all *p* < 0.05 using Fisher’s Exact Test). For these jointly-tuned neurons, the majority exhibited the same preferred task context (i.e. higher firing rate) in all three areas (HPC = 100%, 11/11 neurons, dACC = 88%, 22/25 neurons, preSMA = 65%, 21/32 neurons), which was a greater proportion expected by chance in all areas (p<0.05 using Binomial test).

**Figure 4.**
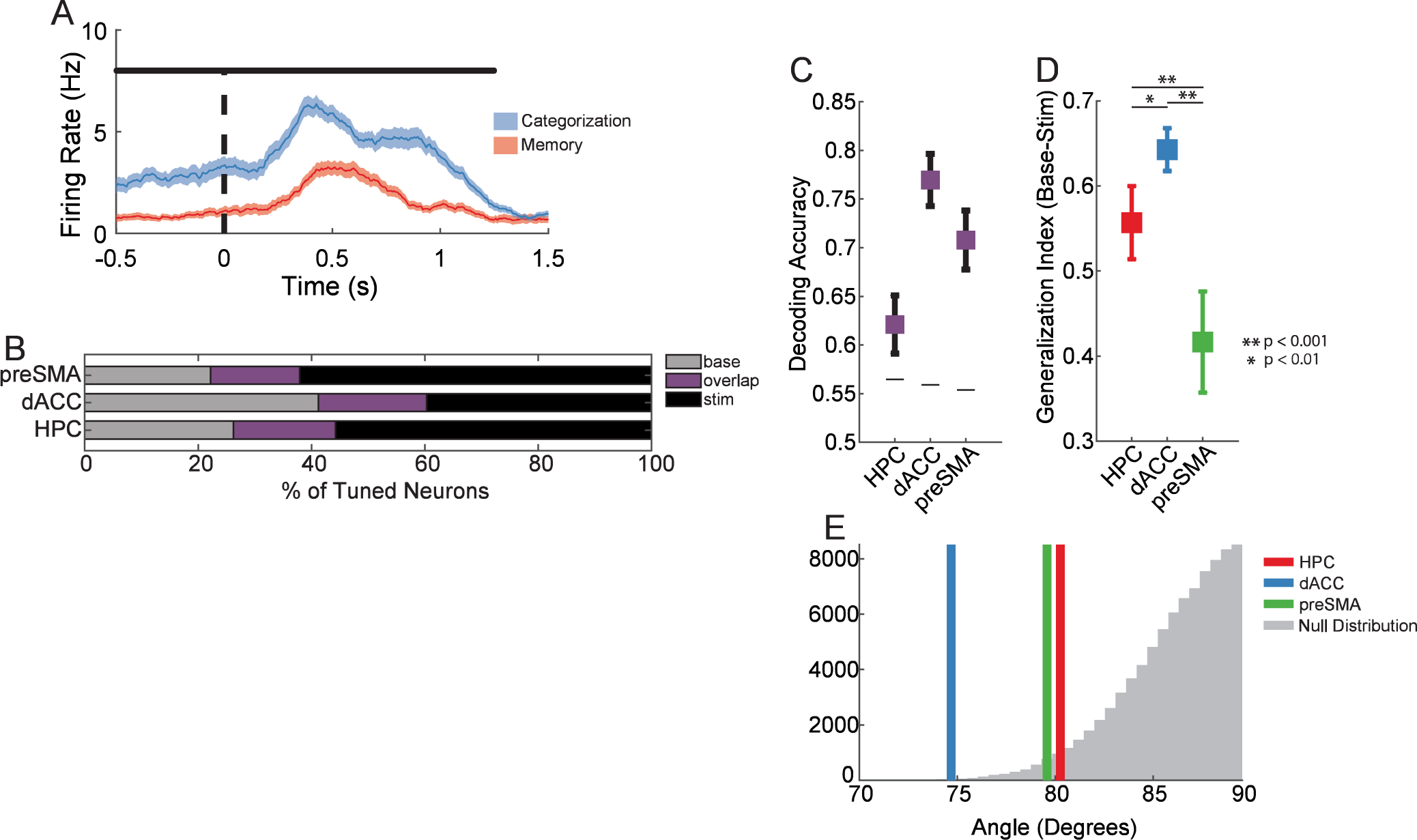
Task context representations generalize between baseline and stimulus periods. **(A)** Example neuron in dACC with context tuning during the baseline and stimulus period. T=0 is stimulus onset. Data points are firing rate mean ± s.e.m. over categorization and memory task trials respectively. Black horizontal line indicates time period where firing rate significantly differs between contexts (1-Way ANOVA, p<0.05). **(B)** Fraction of significantly context-tuned neurons determined using 1-Way ANOVA for context during the baseline (base) and/or stimulus (stim) periods (p<0.05 each). **(C)** Stimulus to. Baseline generalization. Shown is decoding accuracy averaged over both baseline trained/stimulus tested and stimulus trained/baseline tested context decoders. Error bars indicate s.e.m. and horizontal black lines are 95^th^ percentile of null distribution. **(D)** Generalization index for baseline-stimulus context generalization. Values here are computed using the baseline/stimulus context generalization shown in (C), and the within-stimulus and within-baseline context decoding accuracy reported in Fig. 2A. Values reported are mean ± s.e.m. generalization index computed for Hippocampus (red), dorsal Anterior Cingulate Cortex (blue), and pre-Supplementary Motor Area (preSMA). P-values are computed using Wilcoxon Rank-sum test. **(E)** Angles between the baseline and stimulus context-decoding hyperplanes. Plotting conventions identical to Fig. 2C.

We next performed cross-trial period generalization analysis using decoders trained for context between the baseline and stimulus. Mean generalization decoding accuracy, reported as an average over baseline-to-stimulus and stimulus-to-baseline generalization, was significantly above chance for all three regions (Fig. 4C, *p* < 0.001 against Shuffle Null). The generalization index for dACC was significantly greater than HPC (Fig. 4D, blue vs red, *p* < 0.05, permutation test) and preSMA (Fig. 4D, blue vs green, *p* < 0.05, permutation test). This result indicates that dACC exhibited the most temporally stable context representation within the span of a trial. This finding was further confirmed by the angle analysis performed between the baseline and stimulus context decoders, which revealed that only dACC context coding vectors significantly differed from orthogonal (Fig. 4E, blue, *p* = 0.0005, against shuffle null), whereas HPC (Fig. 4E, red, *p* = 0.02, against shuffle null) and preSMA (Fig. 4E, green, *p* = 0.02, against shuffle null) context coding vectors weakly significantly differ from orthogonality when comparing baseline and stimulus. Together, these findings indicate that, while there is some shared neural substrate between the baseline and stimulus context representations in all three areas, the dACC had the representation of context that generalized best on the within-trial timescale among the three brain areas we examined. Note that, in the HPC, while the generalization index was lower than in the dACC, it was still high (0 would be no generalization). This indicates that even in the HPC, task context representations were largely common between these two time periods. Thus, on short time scales, the HPC representation was stable despite it not being stable over the longer time periods of blocks.

### The hippocampal context representation stabilizes under different experimental conditions

Is the temporal stability of context representations in the brain an immutable property intrinsic to each region, or can it vary as a function of experimental setting? We next examined data from Experiment 2 to answer this question. While structurally similar to experiment 1 (compare Fig. 1A with Fig. 5A), context was latent in Experiment 2 rather than explicitly instructed, was defined by arbitrary stimulus-response associations, and needed to be inferred through feedback. Due to these differences, we expect that the processing demands on the hippocampus differ between the two experiments. In experiment 2, patients were rewarded for providing correct binary responses to visual stimuli, with the specific stimulus-response associations learned through trial and error (Fig. 5B). There were two latent contexts, each specified by a different stimulus-response-outcome map. The two contexts alternated at points unknown to the subjects. Thus, subjects had to infer that a context switch had occurred based on the outcomes (error) they received (Fig. 5A,B). No explicit signals were provided to indicate that contexted had switched. Both experiments featured a binary context variable, block structure of similar trial length, a stimulus-identity related variable with four levels, and comparable baseline and stimulus trial periods on which single-neuron and population-level analyses could be performed. These commonalities in the trial-level and block-level structure between experiment 2 and 1 as well as the overlap in recording locations (Fig. S8M) allow for a direct comparison of the encoding strategy employed by the brain at the single-neuron level to represent task-context variables of different kinds (see Methods, Discussion for detailed description of similarities and differences).

**Figure 5.**
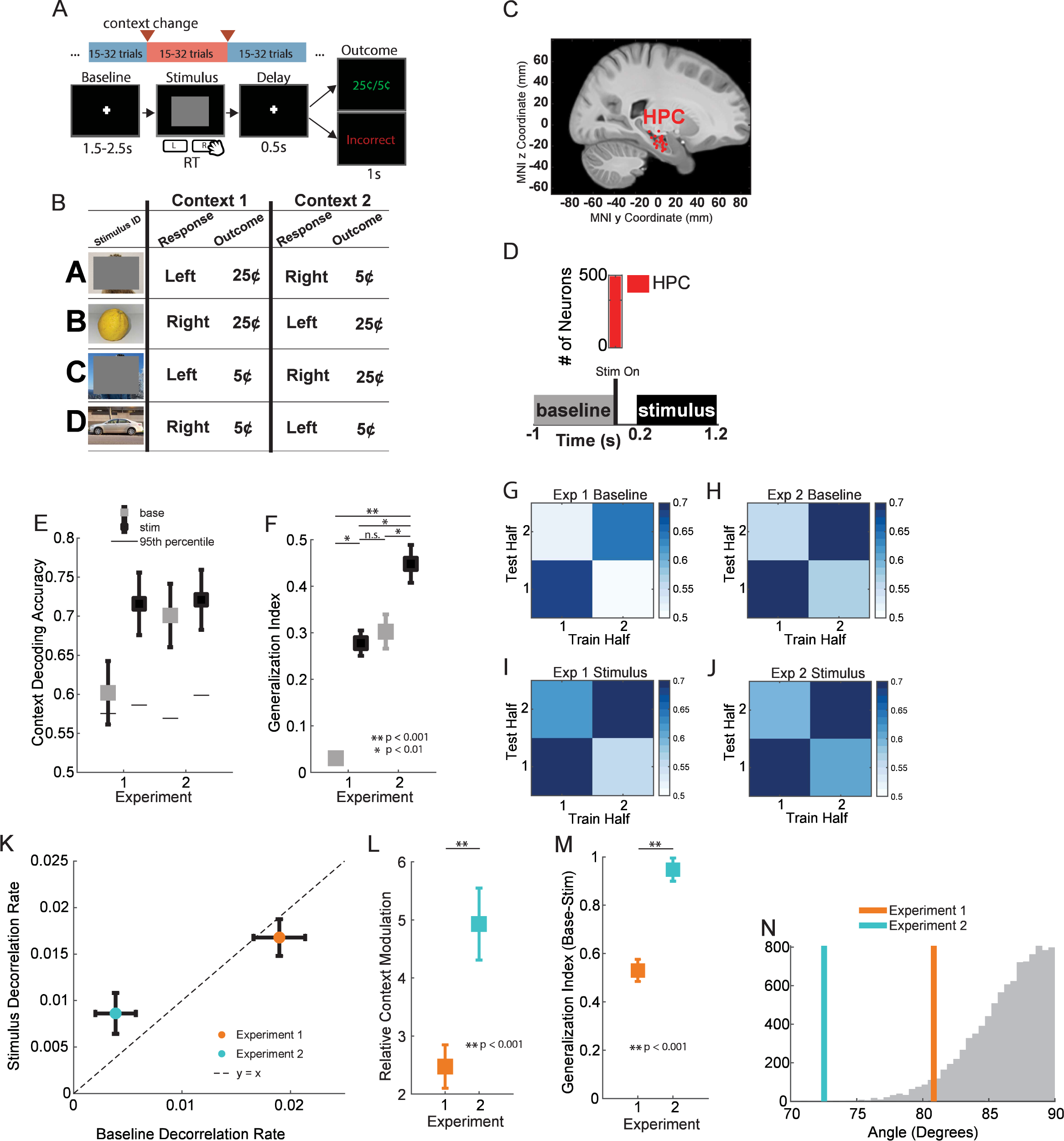
Hippocampal context representation temporally stabilizes when context is latent. **(A)** Experiment 2 consisted of blocks of 15-32 trials where a latent context variable was specified by arbitrary, deterministic stimulus-response-outcome associations. Trials consisted of a pre-stimulus baseline with a central fixation cross, followed by the presentation of a single stimulus (image) to which the patient would respond with a “Left” or “Right” button press according to the current stimulus and context in a speeded manner. Changes in context were covert, but could be inferred from feedback provided during the “outcome” or feedback screen of every trial. Following feedback, the next trial would commence after a jittered delay (1.5-2.5s). **(B)** Electrode locations for the anterior hippocampus (HPC, red). Plotting conventions identical to Fig. 1B. **(C)** Number of single units recorded in the anterior Hippocampus (HPC, red = 499 neurons). **(D)** Context decoding accuracy from HPC during the baseline (gray) and stimulus (black) periods (see inset) using correct trials from both experiments. Black horizontal lines indicate 95^th^ percentile of null distribution. Chance decoding accuracy is 0.5 (two contexts). Values are reported as mean ± s.e.m. **(E)** Cross-temporal context generalization index for both experiments (first half/second half here vs cross-block pair in Fig. 2). See panels (F-I) for illustration. Values reported are mean ± s.e.m. generalization index computed for the baseline (red) and stimulus (blue) periods of each task. P-values are computed by permutation test. **(F-I)** Cross-temporal decoding plots for task context computed across experimental halves instead of across block-pairs are shown for the baseline (F,G) and stimulus (H,I) periods for both experiments. **(J)** Baseline vs Stimulus population decorrelation rate for both experiments. Values are reported as mean ± s.e.m. in each dimension. Dashed line indicates y=x. **(K)** Relative context modulation reported for the HPC during the stimulus period of both experiments. Values are reported as mean ± s.e.m. over iterations of decorrelation curve estimation. P-value is computed by permutation test. **(L)** Baseline-stimulus context generalization index for both experiments. Values are reported as mean ± s.e.m. P-value is computed by permutation test. **(M)** Angles between the baseline and stimulus context-decoding hyperplanes for both experiments. Plotting conventions identical to Fig. 2C.

17 patients completed 42 sessions of experiment 2 (180-320 trials/session, 10-16 blocks/session). Novel stimuli were used in every session. Subjects re-learned stimulus-response-outcome maps in every session. Of these, only sessions where patients exhibited a significant behavioral signature of performing inference on the state of the latent context following covert context switches were considered for analysis. We only considered hippocampal neurons here because our prior work shows that latent context is only represented in the hippocampus (and not the MFC) in this task (see ^12^). Based on these constraints, 325/499 recorded HPC neurons (Fig. 5C,D) from 12/17 patients in 19/42 sessions were included for analysis.

HPC neurons exhibited tuning to context during both the baseline (1-Way ANOVA, p < 0.05 significance, example in Fig. S8A), and to context and stimulus identity during the stimulus period (2-Way ANOVA with interactions, p < 0.05 significance, examples in Fig. S8A,B). The percentage of HPC neurons tuned to context and stimulus identity was not significantly different across the two experiments (Fig. S8C, base context, 10.3 vs 13.2%, p = 0.32, stim context, 22.1 vs 16.3%, p = 0.09, stimulus identity, 17.2 vs 18.5%, p = 0.72, two-tailed Chi-square test). Furthermore, the average ANOVA F-statistic for tuned neurons to context during the baseline or stimulus period was not significantly different between the two experiments (Fig. S8D, *p*_*base*_, *p*_*stim*_ > 0.05, RankSum over neurons). These analyses indicate that single hippocampal neurons exhibited similar univariate tuning properties across the two matched trial periods in the two experiments.

To compare the population-level context code employed by the hippocampus in the two experiments, we performed decoding analysis during the baseline and stimulus periods of both experiments while matching the number of neurons and correct trials per condition across the two experiments. Task context was significantly decodable from the hippocampus in both experiments and time periods (Fig. 5E, *p*_*base*1_ = 0.009, *p*_*stim*1_ = 1.6×10^−6^, *p*_*base*2_ = 9.4×10^−7^, *p*_*stim*2_ = 3.7×10^−5^, using shuffle null distribution). Decodability of context from Exp 1 stimulus, Exp 2 stimulus, and Exp 2 baseline all did not differ significantly from each other (*p*_*stim*1,*base*2_ > 0.05, *p*_*stim*1,*stim*2_ > 0.05, *p*_*base*2,*stim*2_ > 0.05, Permutation Test).

Cross-temporal context decoder generalization (Fig. 5G-J) and subsequent generalization index analysis revealed significantly greater generalization indices in experiment 2 when comparing baseline (Fig. 5F, baseline, gray vs gray, p<0.01, permutation test) and stimulus (Fig. 5F, stimulus, black vs black *p* < 0.01, permutation test) periods across experiments. Notably, cross-temporal generalization indices for context during the stimulus period are greater in experiment 2 despite the fact that univariate tuning to context was greater on average at the single-unit level in experiment 1 (Fig. S8D), and context decoding accuracy did not significantly differ between the two (Fig. 5E, stimulus, black vs black p>0.05, permutation test). These analyses indicate that, from a decoding standpoint, the hippocampal code for context is significantly more temporally stable across blocks in experiment 2 compared to experiment 1 during both stimulus and baseline periods. These effects did not arise from a greater number or more strongly univariately context tuned neurons in experiment 2, and were not driven by differences in recording stability, as the simultaneously encoded hippocampal stimulus representations in both experiments exhibited significant cross-temporal stability throughout experiments (Fig. S8E-G). Furthermore, the stimulus and context coding directions were orthogonal (Fig. S8H), indicating that hippocampus encoded these variables in disentangled subspaces in both experiments.

The increased cross-temporal stability of the hippocampal context representation in experiment 2 suggests that the self-similarity of the neural representation is increased at longer timescales when compared to experiment 1. This prediction was formally tested by performing population-vector autocorrelation analysis on both experiments and comparing both the decorrelation rate and the relative context modulation as was previously performed in experiment 1. These analyses revealed that the hippocampal decorrelation rate was significantly slower in experiment 2 than in experiment 1 during both the stimulus and baseline periods (Fig. 5K, S8I-L). The relative context modulation was also significantly elevated in experiment 2 compared to experiment 1 during both the stimulus (Fig. 5L) and baseline (Fig. S8N) periods. Taken together with the cross-temporal decoding analyses, these findings indicate that the hippocampal neural population was significantly more stable, exhibiting less decorrelation over the timescale of the experiment and maintaining a more stable representation of the task context.

Given the decodability of context during both the stimulus and baseline periods of experiment 2, we next compared the format between the two time periods through baseline/stimulus context decoder generalization analyses. We found that the context baseline-stimulus generalization index was significantly greater for the hippocampus in experiment 2 than in experiment 1 (Fig. 5M, Exp 1 vs Exp 2, *p* = 3.9×10^−4^ RankSum), with coding vectors that deviated significantly from orthogonality (90 deg) for experiment 2 and weakly in experiment 1 (Fig. 5N, angle vs. chance *p*_*Exp* 1_ = 0.036, *p*_*Exp* 2_ = 3×10^−4^). These findings suggest that the persistent representation of context generalizes across time periods within an individual trial in experiment 2. Thus, taken together, these analyses indicate that the latent context variable in experiment 2 is encoded in in a more temporally stable manner simultaneously at the timescale of a single trial (∼2s) and at the timescale of the experiment (∼20min) in the hippocampal representation.

## Discussion

We found that neural populations in the human medial frontal cortex form a temporally stable representation of instructed task context that persists over many minutes in the absence of re-cuing both during baseline periods and stimulus processing periods (Fig. 2). In contrast, in the hippocampus, representations of instructed task context were encoded dynamically over the same long time periods. The changes in task context encoding occurred rapidly at task boundaries with stable coding within blocks (Fig. S4). While neurons in all areas were found to exhibit slow decorrelation (Fig. 3), this effect was slower in MFC. Decorrelation alone did not explain the effects of task context, because task context caused abrupt changes at the point of task switching. Lastly, we found that in a different experiment with similar high-level task structure but with radically different trial-level demands, hippocampal context representation became significantly more stable across time (Fig. 5), indicating task dependence.

### Medial frontal cortical neurons representing task context

Medial frontal cortical structures in the primate brain have long been appreciated for their role in maintaining representations of task context variables that support persistent behaviors. However, previous single-neuron studies in non-human primates frequently provide task context cues on a trial-by-trial basis, thus obviating the need to maintain the instructed variable beyond a single trial. In contrast, here subjects needed to maintain an instructed task context over many trials over many minutes. In experiment 1, in both dACC and preSMA, task context was represented stably across time with high cross-temporal stability, slow rate of population-level decorrelation over time within-context, and persistent encoding of task context by individual neurons (Fig. 3H,I). This gave rise to the ‘checkerboard’ pattern of cross-correlation seen in Fig. 3. There are clear computational advantages for a network to employ such a static context representation, most notably the ability of a downstream region to read out the current task context arbitrarily long after the cue has been provided. Thus, the presence of medial frontal cortical context representations that generalizes across long and variable time periods could facilitate the ability of the individual to flexibly maintain persistent behavior. For context, recurrent neural network and transformer-based network^40,41^ approaches typically struggle with generalization to sequence lengths outside of their training distribution. This flaw in artificial neural networks could possibly be ameliorated by encouraging the learning of temporally disentangled representations similar to those we observe in the MFC.

What mechanism could allow for a neural code of task context to be persistently maintained for such long periods of time? Assuming a spike train autocorrelation time constant τ_s_ = 350msec for MFC^42^, the ratio to the average block duration B/τ_s_, during which the context representation was stable, is ∼ 300 (450 for the control variant) in experiment 1. Various circuit-level mechanisms, including recurrent excitation and short-term synaptic plasticity^43,44^, do provide potential explanations for the increased window of temporal integration exhibited by primate frontal cortical neurons, and account for both static and dynamic codes those neurons exhibit. However, these models and analyses are typically limited to the duration of single-trials in working memory tasks, and representations of task context variables do not need to be maintained for more than 2-3 seconds during those delays. Neurons in some models can exhibit long time constants (up to 4 seconds in Area 24)^45^, but it is unclear if such models can explain task variable coding activity that persists 2 orders of magnitude longer. Our findings call for future work on neural circuit-level models for maintaining information with persistent activity for up to minutes.

One notable aspect of our analysis of the MFC is that dACC and preSMA neurons differed with respect to baseline to stimulus generalization. dACC neurons exhibiting considerably greater context coding direction alignment between the two trial periods when compared to preSMA. These findings support the role of the dACC as a temporal storage buffer for context variables that influence behavior in a temporally extended way, since one might expect that a storage buffer would need to stably encode the variable it has buffered to facilitate flexible readout. Compared to the preSMA, the difference in temporal stability of the context code across trial phases could result from intrinsic differences in spike train autocorrelation of neurons in these regions, which are longer in the Anterior Cingulate Cortex when compared to more caudal frontal cortical regions^42,44,46^.

### The logic underlying static and dynamic hippocampal codes for task variables

The cognitive map formed by hippocampal neurons has been extensively studied for its encoding of a wide variety of variables in support of flexible behavior. Here, we considered variables that can be split into two different categories: task context variables, which determine the appropriate response for many different stimuli, and stimulus variables, which encode identity or category information about a currently presented visual stimulus. The task context variables in these experiments also differed from stimulus variables in that their value needed to be remembered for minutes at a time. We found that the code for task context in experiment 1 was dynamic in HPC and static in MFC.

What might be the reason for a dynamic code in the hippocampus? Several potential interpretations come to mind. First, it is possible that the presence of static MFC task context codes could obviate the need for temporal stability in the HPC context representation. If a static code persists elsewhere, the hippocampus is free to return to its “default” state of internally generated cell assembly sequences, which do not cross-temporally generalize^34^. This explanation could also account for the increased cross-temporal stability in the hippocampal context code observed in experiment 2, in which frontal cortical context representations were largely absent^12^. Second, task context was signaled explicitly in experiment 1 but had to be inferred from feedback in experiment 2. The hippocampus supports inference behavior^11,47–49^, and bilateral temporal lobectomy patients are unable to perform tasks with inferred rules such as the Wisconsin Card Sorting Task^50^ (which is similar to experiment 2). Thus, a potential interpretation of our data is that the hippocampal context representation may stabilize across time specifically when it is needed to support persistent behavior, i.e. when the task context variables are latent and must be inferred and are not persistently encoded elsewhere in the brain.

The hippocampus plays a prominent role in memory formation, and the presence of episodic memory demands in experiment 1 (one of the two tasks was a recognition memory task) could create a demand for the hippocampus to encode the passage of time, i.e. the current temporal context, alongside the instructed contexts in experiment 1. In experiment 1, the tuning of single hippocampal neurons to task context was better accounted for by block-specific tuning, and as a population the neurons exhibited continual decorrelation on the timescale of minutes that was comparable in magnitude to the effect of switching tasks between blocks (hippocampal RCM = 3.2). The encoding of time in the human hippocampus is achieved at the single-neuron level through time cells and ramp cells^33^, and at the population level by sequential firing of neurons. Supporting this view, temporal context is reinstated in the human hippocampus during successful retrieval^37^. In our data, the hippocampus multiplexed temporal context information with task context information such that both variables were simultaneously encoded in the same neural population, possibly reflecting the association of multiple behaviorally relevant high-level context variables^51^. Such temporal context encoding may have been absent from experiment 2 due to the small number of stimulus-response associations to be remembered and the lack of behavioral demand to form and/or retrieve an episodic memory. New experiments are needed to specifically test this hypothesis.

What strategies are employed by the hippocampus for organizing the simultaneous representation of task context variables and stimulus variables in the same neural state space? Evidence from modern systems neuroscience points towards the hippocampus simultaneously exhibiting high and low dimensional properties in its state space representation of the environment, where combinations of these variables are mixed to varying degrees, allowing for flexible readout for many downstream tasks while retaining some advantageous geometric properties that allow for generalization of one variable across others^11,12,52,53^. Here, we have demonstrated in two different experiments that hippocampal representations of stimulus variables are orthogonally encoded with respect to the blocked task context variables and the passage of time. Could the segregation of stimulus and context variables into orthogonal subspaces be a general feature of hippocampal representations? Place-field-like coding across conjunctions of variables is frequently seen in the hippocampal representations of rodents and non-human primates, arguing against this segregation as a general property of the hippocampus^54–56^. However, this may strictly be a property of human hippocampal representations, and the disentangling of stimulus and context codes may underlie the rapid learning and generalization behaviors exhibited by humans when compared to other species^11,12,57^.

## Methods

### Participants

The study participants were adult patients being surgically evaluated for invasive treatment of drug-resistant epilepsy (see Table 1). These patients were treated at Cedars-Sinai Medical Center (CSMC) and Toronto Western Hospital (TWH). All patients provided informed consent and subsequently volunteered to participate in this study. All research protocols were reviewed and approved by the institutional review boards of CSMC, TWH, and the California Institute of Technology.

### Experiment 1

One group of 13 patients (see Table 1) performed 33 sessions an experiment that required alternation between two task contexts, one requiring semantic categorization of visual stimuli (categorization task), and the other requiring recognition memory of those same stimuli (memory task). Patients performed eight blocks of 40 trials, with the required task alternating between categorization and memory across consecutive blocks. Patients always began with categorization for the first block of the session. Text-based instructions for the required task in the current block were provided at the start of each block, and were not re-cued until the next block thus requiring patients to persistently remember the current task being executed. Patients could spend as much time as needed on instruction screens, and voluntarily proceeded into every new block after reading the instructions. Both tasks were formulated as binary (yes/no) questions of the form: “Is this an image of an X?” for the categorization task, where X was one of four unique semantic categories, and “Have you seen this image before?” for the memory task. The number of new and old images of each semantic category were balanced in each block to prevent response biases and to facilitate balanced decoding analyses. Individual trials consisted of a jittered pre-stimulus baseline (1s to 2s) followed by image presentation for a variable amount of time until the patient’s response for that trial was provided. Patients provided trial responses using either a left/right button press on a CEDRUS binary response box or by saccade left/right to indicate True/False for each trial. The response modality was randomized over blocks and was re-cued every 20 trials. Following the response, the stimulus was removed from the screen and the baseline period for the next trial was initiated. Trial-by-trial feedback was not provided.

### Experiment 1 Control Variant

A control variant of the experiment described above, which was designed to disentangle stimulus processing from decision variables and motor plans, was utilized for a fraction of the sessions (5/13 patients, 6/33 sessions). In this variant, instead of being allowed to respond freely as soon as the image appeared on the screen, images were presented for a fixed interval (1s), then trial responses were allowed after a jittered delay (0.5s to 1.5s) when a response cue appeared on the screen. These task variations were uniformly applied to all trials, and thus could not have generated a bias in the representation of one task condition over another. Furthermore, none of the analyses shown here were performed on a response-aligned time window. Nevertheless, the core analyses in this work were re-conducted on neural pseudopopulations constructed exclusively from these control sessions, and all findings were re-capitulated (See Fig. S4).

### Experiment 2

A separate group of patients 17 patients (see Table 1) performed 42 sessions of a second experiment (180-320 trials/session, 10-16 blocks/session) that shared key structural elements with the first experiment both at the trial level and at the block level that allowed for direct comparison of neurophysiological task responses across the two experiments. In sessions of this experiment, patients learned arbitrary stimulus-response-outcome (SRO) associations for four unique stimuli arbitrarily associated with either a left or right button press in one of two latent contexts. The contexts were related in that the required response for each stimulus was inverted between the two contexts (e.g. stimulus A was associated with left button press in context 1 and right button press in context 2, etc..). Blocks consisted of 15-32 trials in a given context before a covert switch to the other context. Trials consisted of a pre-stimulus baseline (1.5s to 2.5s), followed by a speeded response (left/right button press) provided with the onset of the stimulus, and the presentation of an outcome (either reward or “incorrect”) for 1s following a fixed 0.5s delay. Rewards were provided deterministically such that, if a patient had learned the SRO map in a given context and suddenly encountered an incorrect trial, this was an unambiguous signal that the state of the latent context variable had changed. Patients could learn to perform inference on the state of the latent context variable such that, after a single incorrect trial, patients could infer that the context had changed and update all stimulus-response associations in accordance with the new context.

### Cross-Experiment Comparison

The two experiments considered here share a considerable amount of structure that facilitates cross-task comparison. Both experiments contained a binary task context variable that was designed to elicit different responses for the same stimuli depending on the state of the context variable. Both experiments have a blocked structure such that the context variable varies slowly with time, and many trials must be completed in a given block before the context changes. The current context is not re-cued in either experiment, with explicit instructions being provided once at the beginning of each block in experiment 1 and never being provided in experiment 2, thus requiring a persistent representation of task context in both experiments to achieve high performance. Trial structure is also very similar between the two experiments. In both cases, trials consist of a pre-stimulus baseline where a single gray fixation cross is present on the screen for ∼2s. Trial onset is marked by the appearance of a single image subtending ∼10 visual degrees. The image is removed when the patient provides a response for that stimulus in accordance with the currently instated context. Responses were formulated as binary in both experiments for all task contexts, with the Categorization and Memory tasks in experiment 1 formulated as yes/no questions and the two latent contexts in experiment 2 requiring left/right button presses. Thus, for the time periods analyzed here including baseline (-1s to 0s prior to stimulus onset) and stimulus processing (0.2s to 1.2s following stimulus onset), patients were engaged in cognitive tasks with roughly similar structure and comparable cognitive demands. Elaboration of the differences between these experiments and associated limitations is provided in the discussion.

### Electrophysiology

#### Electrode Placement and Recording

Extracellular electrophysiological recordings were conducted using microwires embedded within hybrid depth-electrodes (AdTech Medical Inc.) implanted bilaterally into the hippocampus, amygdala, dorsal anterior cingulate cortex, pre-supplementary motor area, ventromedial prefrontal cortex, in addition to variable unilateral or bilateral electrodes in ventral temporal cortex as determined by clinical needs. Broadband potentials (0.1Hz – 9kHz) were recorded continuously from every microwire at a sampling rate of 32kHz (ATLAS system, Neuralynx Inc.). All subjects included in the study exhibited voltage waveforms consistent with well-isolated single-neuron action potentials in at least one implanted microwire.

#### Electrode Localization

Electrode localization was conducted using a combination of pre-operative MRI and post-operative CT using standard alignment procedures as previously described^6^. Electrode locations were also co-registered to the to the MNI152-aligned CIT168 probabilistic atlas^58^ for standardized location reporting and visualization. Placement of electrodes in gray matter was confirmed through visual inspection of subject-specific CT/MRI alignment, and not through visualization on the atlas.

### Spike Detection and Sorting

Raw electric potentials were filtered with a zero-phase lag filter with a 300Hz-3kHz passband. Spikes were detected and sorted using the OSort software package^59^. All spike sorting outcomes were manually inspected and putative single-units were isolated and used in all subsequent analyses. All processing and analysis of neural data was performed using MATLAB (The Mathworks, Inc., Natick, MA).

### Analysis Periods, Single-Neuron Tuning, and Construction of Pseudo-populations

All analyses were conducted on firing rates of neurons computed during two trial epochs the baseline period (base), defined as -1s to 0s preceding stimulus onset on each trial, and the stimulus period (stim), defined as 0.2s to 1.2s following stimulus onset on each trial. Firing rate vectors for every neuron were constructed during both trial periods.

Single-neuron tuning properties were assessed using univariate and multivariate ANOVAs applied to the firing rate vectors for each neuron independently unless otherwise stated. Task context was encoded as a categorical variable with two levels for both experiments. Semantic image category in experiment 1 and stimulus identity in experiment 2 were encoded as categorical variables with four levels. Any references to depth-of-tuning of a neuron for one of these variables or their interaction (e.g. Context x Stimulus identity) refer to the F-value of the variable in question when the ANOVA is performed on the trial-level firing rate vectors either during the stimulus or baseline periods. Note: all ANOVA analyses were performed using spikes counted during correct trials for the stimulus period and during baselines preceding correct trials. For some control analyses, an ANOVA F-statistics distribution matching procedure is also performed between neurons recorded in different regions. To match F-statistic distributions, valid pairs of neurons were identified, one from each region, whose F-statistics for the context variable were within 0.1. The candidate pair was removed from the pool of available neurons, and another pair was selected until no more valid pairs were present, at which time all remaining neurons were excluded from the subsequent distribution-matched analysis. This procedure creates two populations of neurons, one for each region participating in the balancing procedure, whose ANOVA F-statistic distributions are statistically indistinguishable.

Firing rate vectors including all trials for single neurons were concatenated to create neural pseudo-population matrices of dimension (# of trials x # of neurons) on which all subsequent decoding analyses were performed. These pseudo-populations only consisted of neurons that exhibited at least 0.1Hz firing rate averaged over the entire recording session. Repeated recording sessions of a given experiment with a given subject were typically separated by several days, and neurons recorded during these repeated sessions were treated as independent neurons in the pseudopopulation. No stimuli or stimulus-response pairings were ever re-used in repeated recording sessions for either experiment, thus preventing potential behavioral and neural confounds related to recognition memory signals across recording sessions.

### Trial-Balanced Decoding Analysis

Decoding analyses were performed using a linear support vector machine, and all decoding accuracies are reported out-of-sample using 5-fold cross-validation unless otherwise specified (e.g. cross-condition generalization). Model fitting was performed using the “templateSVM” with a linear kernel and the “fitecoc” model-fitting methods from the Stats toolbox of Matlab 2021b. Trial-balanced decoding of a task variable was conducted by concatenating firing rate vectors for neurons in a given region to construct firing rate matrices, i.e. the pseudopopulation response matrix, with dimensions KT x N, where K is the number of conditions (typically 2 apart from image category decoding, where there were 4 categories), T is the number of correct trials per condition, and N is the number of neurons. Neurons were excluded from the pseudopopulation if they there were fewer than 15 correct trials per condition in that recorded session. Neurons with more than the minimum number of correct trials had their trials randomly sub-sampled so that the number of correct trials per feature and per condition could be matched prior to decoding. To account for the presence of noise correlations between simultaneously recorded neurons, trials were also shuffled independently for each neuron within-condition prior to decoder fitting. To account sub-sampling and randomization bias, the trial sub-sampling and within-condition shuffling procedure were repeated 250 times, with reported decoding accuracies being the average over these repeats. All error bars shown are standard deviations over the distribution of decoding accuracies unless otherwise specified. Null distributions for decoding analyses were constructed by shuffling condition labels and reporting out-of-sample decoding performance, again with 250 repetitions. The significance of individual decoding accuracies was determined by reporting the p-value of the average decoding accuracy against the gaussian maximum likelihood fit of the null distribution. The significance of the difference between decoding accuracies (e.g. between two areas) was determined by reporting the p-value of the true difference against a null distribution of the difference constructed by computing all pair-wise differences between points in the null distributions for each of the two decoding accuracies being considered.

### Decoder Cross-Condition Generalization

Generalization analyses for decoders are performed by training a decoder to discriminate between two states of a variable in one condition and testing whether that decoder performs above chance in discriminating the same variable states in another condition. Critically, in order for such analysis to be performed, the variable in question must un-ambiguously be specified in the source condition (on which training is performed), and the target conditions (on which testing, or generalization is performed). In the case of task context, this generalization performance was performed over trial phases (baseline/stimulus), block phases (first block half/second block half), and over experiment phases (block pairs). Furthermore, generalization analysis requires that the training and testing feature spaces are aligned. In this case, the requirement of a minimum number of 15 correct trials per neuron per condition was also independently applied to the source and target conditions for these generalization analyses. For example, if training a context decoder on blocks 1/2 and testing on blocks 3/4, a neuron must have at least 15 correct trials per context in blocks 1/2 and 3/4 independently. Neurons that did not meet these criteria separately for both source and target conditions were excluded from the analysis. Since in a generalization analysis, decoding accuracy is out-of-sample by construction, cross-validation was not used, and all available trials were used for training and testing. However, since trial sub-sampling and within-condition shuffling were also employed here, generalization decoding accuracies were also reported as the average over 250 repetitions.

To control for the amount of task variable information available to a decoder during training, a generalization index is reported that normalizes the performance of the decoder in the generalized conditions to the out-of-sample performance of the decoder in the training conditions. Specifically, the generalization index is computed as:

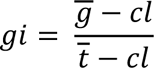

where *ḡ* = mean performance over all instances of generalization for all trained decoders, *t̄* = mean out-of-sample training performance for all decoders, *cl* = chance level. For example, when computing the generalization index for context decoders over block pairs, for the 8 blocks, *t̄* is the average over training performance of 4 context decoders, *ḡ* is the average performance over 12 instances of generalization (note: generalization performance of decoders is not necessarily symmetric), and chance level is 0.5.

### Coding Vector Angles

The coding vectors used in all angle analyses are the β coefficients of decoders trained to decode task variables, typically task context, in different phases of each experiment. These coefficients are the weights returned by the SVM model fitting procedure. They reflect the relative contribution of each feature (neuron) to the decoder, with better-tuned neurons to the variable in question being assigned higher magnitude coefficients. The coefficients are also signed to reflect which of the conditions that feature prefers (e.g. the context assigned to +1 or -1 for binary classification). Class label assignment for classification was kept constant to allow for decoder generalization and meaningful estimation of the angle between coding vectors for different decoders. Angles between coding vectors were computed in by applying the definition of the dot product in N-dimensional neural state spaces, where N was the number of neurons included in the given analysis. N was matched for all regions within a given analysis so that angles reported in the same plot were directly comparable, and not computed using vectors with different dimensions. All angles between coding vectors were reported as the average over the 250 repetitions of decoder estimation described above. Null distributions were constructed by pooling together all possible pair-wise angles between shuffle-null decoders trained as described in the “Trial-Balanced Decoding Analysis” section. Angles are computed between: context decoders trained on different block pairs, context decoders trained on the baseline and stimulus processing periods, and image category decoders trained on different block pairs. In all these cases, the same neurons are used as the features in the two decoders between which the angle is being computed, so the neural state spaces are aligned by construction and the angle between coding vectors is readily interpretable as an overlap in the coding direction for the variable being decoded.

### Population Vector Autocorrelation

The trial-level autocorrelation of the neural population in each region was estimated computing the Pearson correlation between population vectors for every pair of trials present in each experiment. For experiment 1, all trials were included in this analysis leading to 320^2^ dimensional autocorrelation matrices. Population vectors here were sub-sampled to match the smallest number of neurons available in any region as previously described so that correlation values were directly comparable across regions. For experiment 2, since block lengths were randomized for every block in every session, all blocks were sub-sampled in length to match the smallest available number of trials in a given block. Re-sampled estimation of population vector autocorrelation for this experiment was simultaneously performed over neurons and trials-in-block. Reported autocorrelation heat maps are an average over 250 repetitions of re-sampled estimation, and are convolved with a 2D Gaussian filter with a standard deviation of 1 for visualization purposes. All subsequent analysis on population autocorrelations was performed on the un-smoothed maps. Block-wise decorrelation curves are computed by taking the average pairwise correlation between all trials within the same block for block distance 0, average pairwise correlation between all trials one block apart for block distance 1, and so on. On-diagonal correlations (trial with itself) are ignored to prevent artificial inflation of block distance – correlation. Even and odd block distances are colored differently to reflect the fact that even block distances correspond to trial-level correlations within the same task context and odd block distances correspond to trial-level correlations between different task contexts, since task contexts alternated at the block level in both experiments.

Two metrics are further derived from these curves: the decorrelation rate and the relative context modulation. The decorrelation rate for the neural population in each region was quantified by performing linear regression on the decorrelation curves and reporting the absolute value of the estimated slope. This slope was always negative for all neural populations and time periods considered as there was no instance in which the self-similarity of a neural population increased over time. The decorrelation rate is reported in units of *block*^−1^since the slope is an estimate of the change in linear correlation of the population (unitless) divided by the block distance, which is measured in units of blocks by definition. The relative context modulation is defined as the average absolute difference in linear correlation between trials 0 blocks apart and trials 1 block apart, normalized by the decorrelation rate. Since the absolute block 0-1 difference is also computed in units of *block*^−1^, the relative context modulation is a unitless quantity that reports the effect of re-cuing task context on a population of neurons, normalized to the baseline tendency for that neural population to decorrelate over time. A relative context modulation of 1 indicates that the change in representation experienced by a neural population over the timespan of a block due to intrinsic decorrelation and due to explicit cueing of a different task are equivalent, with values greater than 1 and less than 1 indicating dominance of task-recuing effects and intrinsic decorrelation respectively. Reported values for both the decorrelation rate and the relative context modulation are averages over the 250 repetitions of re-sampled autocorrelation estimation, and error bars are s.e.m. over these repetitions.

## Acknowledgments.

We thank R. Adolphs for advice and support throughout all stages of the project, and J. Minxha, M. Meister, R. Andersen, and S. Fusi for discussion. We thank all subjects and their families for their participation and the staff and physicians of the Cedars-Sinai and Toronto Western Epilepsy Monitoring Units for their support. This work was supported by the BRAIN Initiative through the NIH Office of the Director (U01NS117839 to U.R.), the Simons Foundation Collaboration on the Global Brain (to U.R.), the Moonshot R&D JPMJMS2294 (to Kenji Matsumoto), and by a merit scholarship from the Josephine De Karman Fellowship Trust (to H.S.C.).

## Author contributions

Conceptualization: H.S.C, U.R., Data analysis: H.S.C., Writing: H.S.C., U.R., R.A. Surgeries: A.N.M. and T.A.V., Supervision: U.R., R.A., and A.N.M.

## Data availability statement

Data used for both experiments is publicly available, see XXX.

---

**Figure S1.**
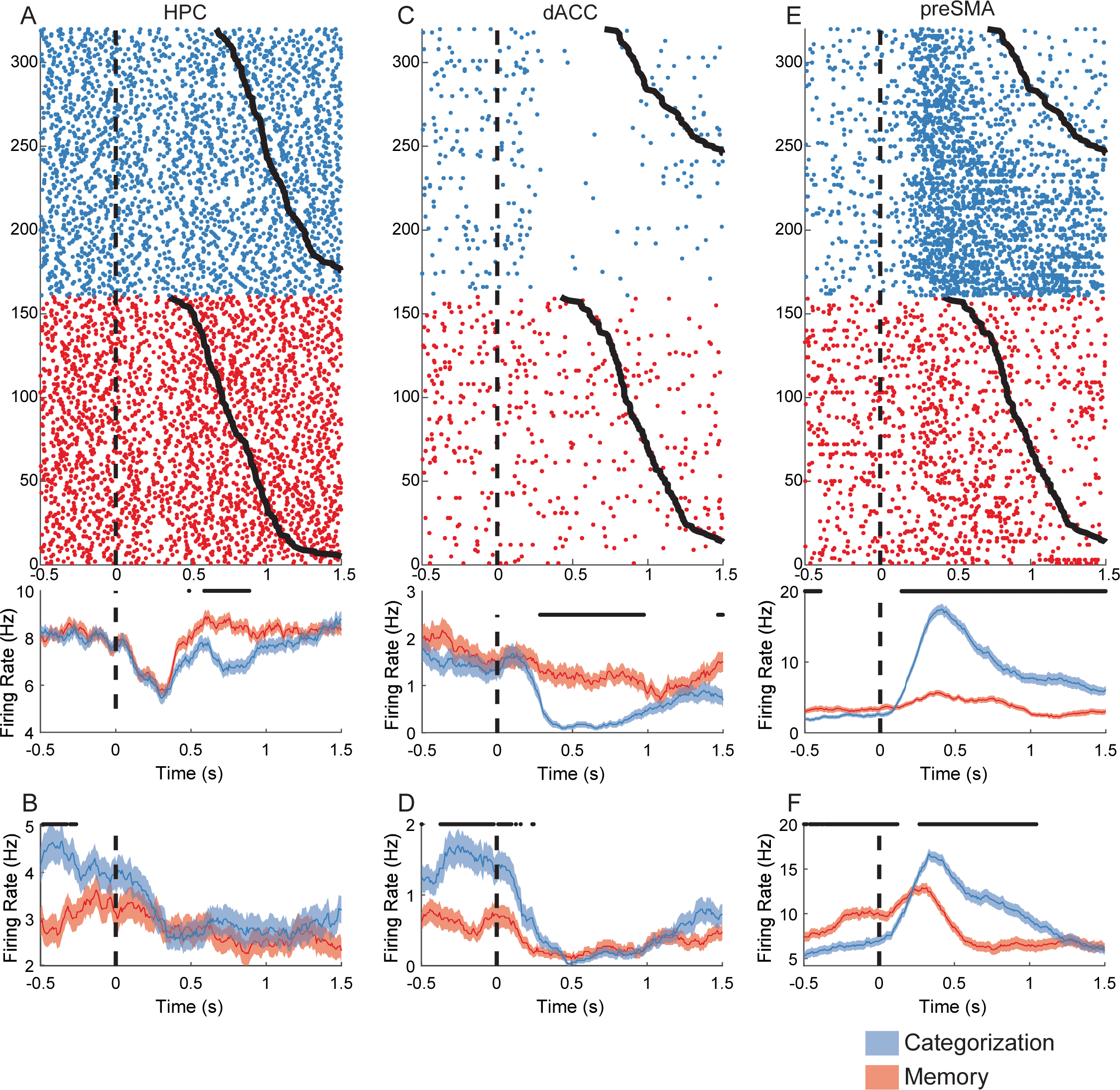
Example neurons recorded in Task 1 that exhibit context tuning during stimulus and baseline periods. **(A)** Example raster (above) and PSTH (below) for a neuron in anterior hippocampus (HPC) that was context-tuned during the stimulus presentation period. Trials in the raster are re-ordered according to task context (categorization = blue, memory = red), and are sorted according to reaction time therein, as indicated by the black curves on the right. Vertical dashed line denotes stimulus onset. PSTH shows mean ± s.e.m. firing rate computed over trials. The black dots above the plot indicate time periods where firing rate significantly differs between contexts (1-Way ANOVA, p<0.05). **(B)** PSTH shown for a different neuron in HPC that exhibited significant context tuning during the baseline period prior to stimulus onset (i.e. to the left of the vertical dashed line). **(C,D)** Same as **(A,B)**, but for dorsal Anterior Cingulate Cortex (dACC). **(E,F)** Same as **(C,D)**, but for pre-Supplementary Motor Area (preSMA).

**Figure S2.**
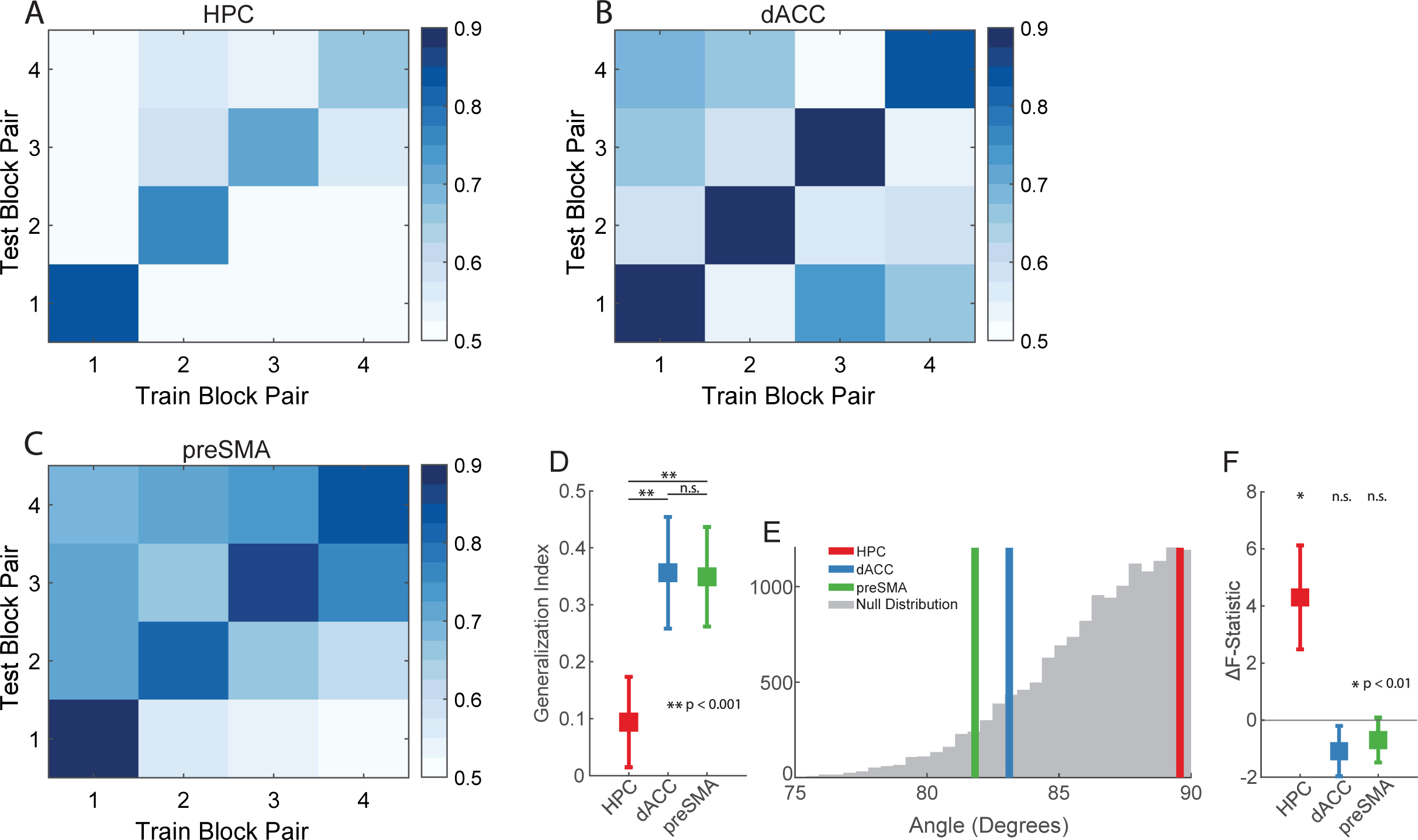
Temporal generalization of task context representation during the baseline period. **(A-C)** Cross-temporal decoding plots for task context computed during the baseline period (-1s to 0s prior to stimulus onset). X-axis indicates which block pairs are used to train the context decoder, and y-axis indicates the block pairs on which the decoder is evaluated. Plots are shown for HPC **(A)**, dACC **(B)**, preSMA **(C)**. Plotting conventions identical to those in Fig. 2B-D. **(D)** Generalization index computed during the baseline for the cross-temporal generalization of context decoding across block pairs. Plotting conventions identical to those in Fig. 2E. P-values are computed by permutation test. **(E)** Angles computed between vectors normal to the hyperplanes of the baseline block-pair context decoders. Plotting conventions identical to those in Fig. 2F. **(F)** Single-unit model comparison of ANOVA F-statistics for block number vs task context. Plotting conventions identical to those in Fig. 2G.

**Figure S3.**
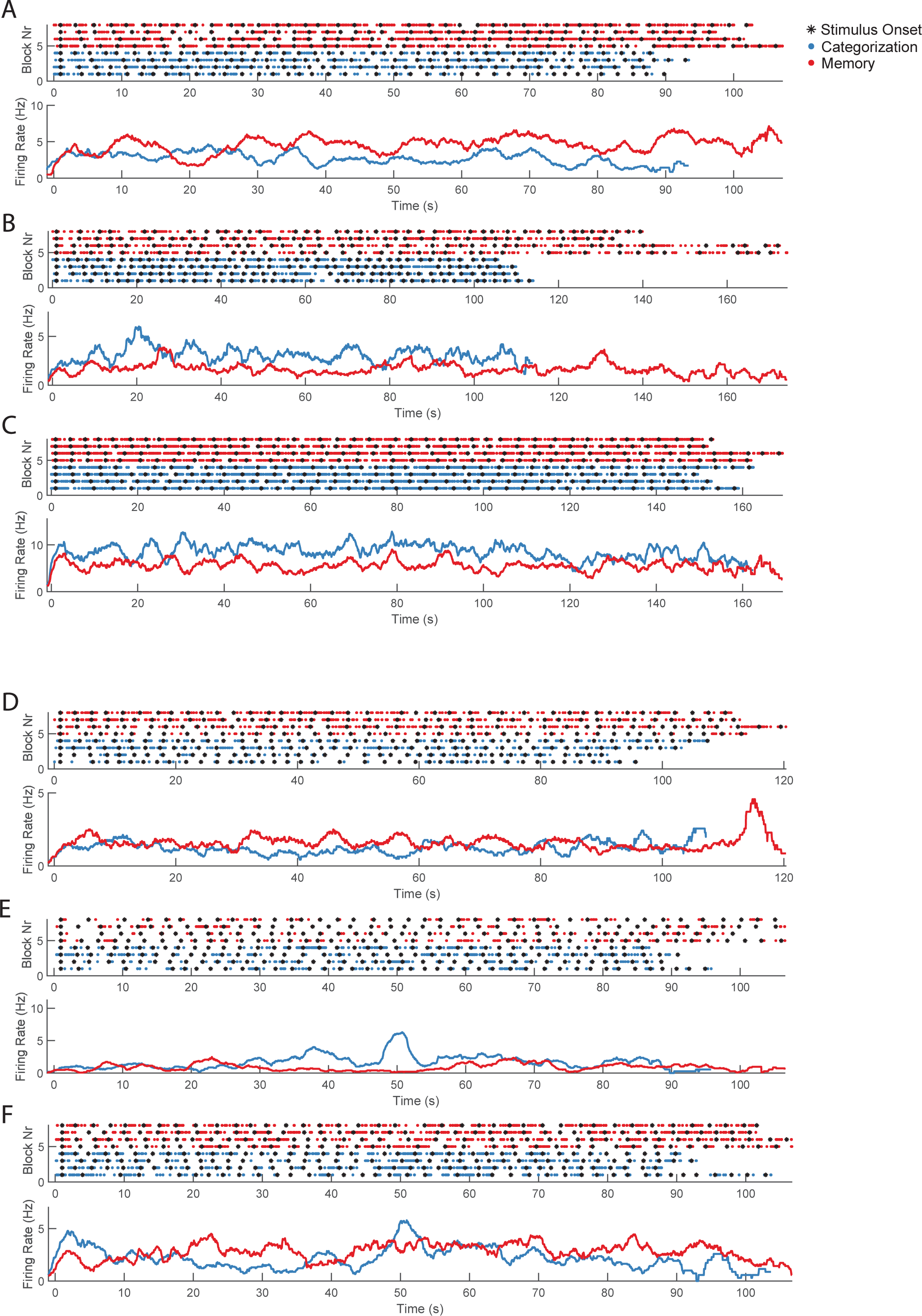
Single-unit rasters/PSTHs showing persistent activity over entire blocks. **(A)** Example raster (above) and PSTH (below) for a neuron in the dorsal Anterior Cingulate Cortex (dACC) that exhibited persistent firing rate context modulation throughout entire blocks. An individual row in the raster (above) corresponds to the activity of a single neuron plotted for a block. Each point corresponds to one spike. Black stars indicate stimulus onset times. Blocks are re-ordered according to task context (categorization = blue, memory = red), and are aligned to the stimulus onset time of the first trial in each block. PSTH (below) shows mean firing rate computed over blocks. Since block durations differ, due to randomization of inter-trial intervals and variability in patient responses, a blocks ceases to contribute to PSTH after the final spike in that block is discharged. **(B-C)** Same as **(A)**, but for pre-SMA. **(D-F)** Same as **(A)**, but for HPC.

**Figure S4.**
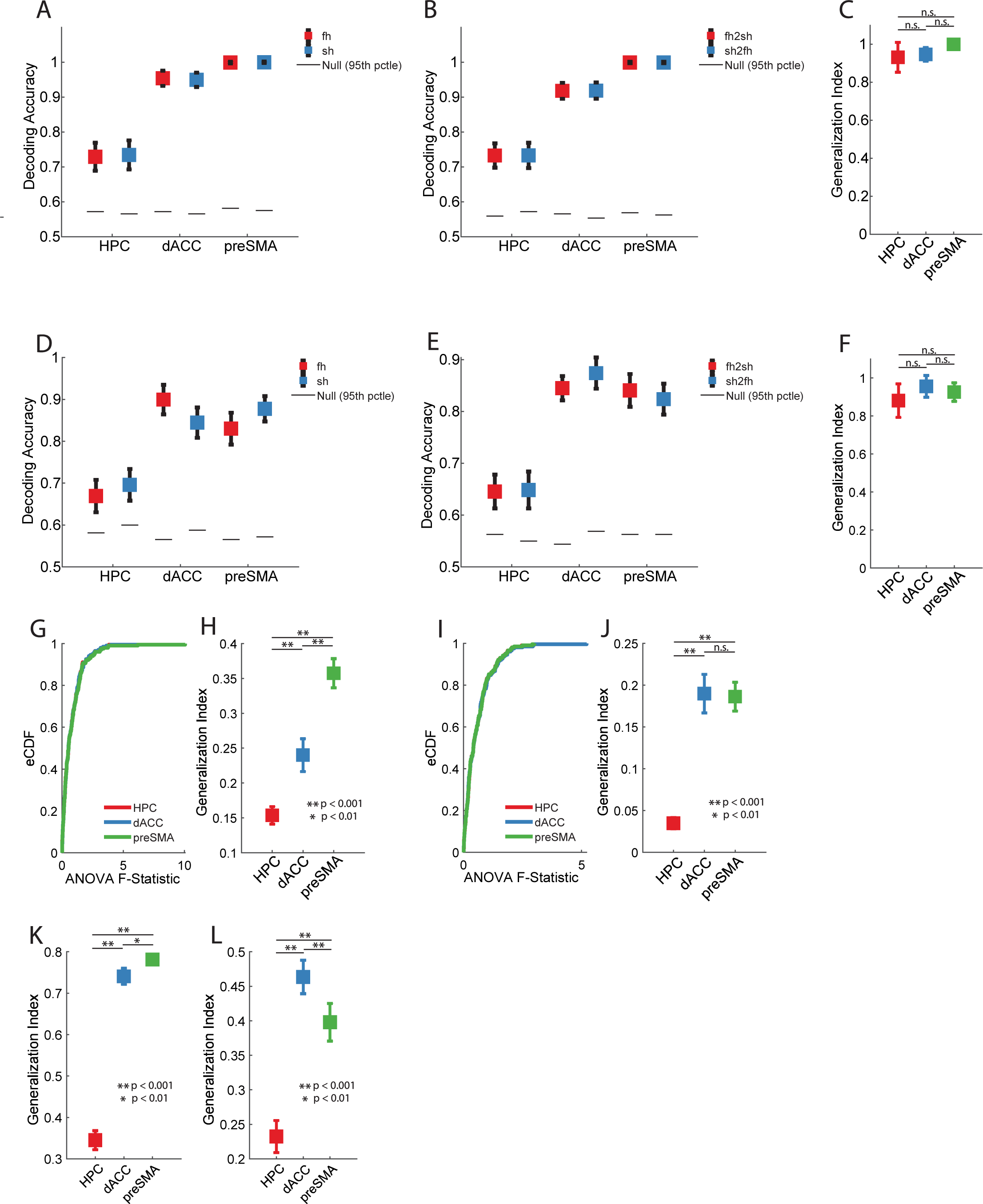
Control analyses for temporal generalization of context representation. **(A)** Task context decoding during the stimulus processing period using the first-half (fh, red) and second-half (sh, blue) of every block pair to demonstrate within-block context decoding stability. Plots show mean decoding accuracy ± s.e.m. over bootstrap iterations. Horizontal black lines indicate 95^th^ percentile of shuffle null. **(B)** Same as **(A)**, but for generalization decoding accuracy of the context decoder from the first block half to the second block half (fh2sh, red) and from the second block half to the first block half (sh2fh, blue). **(C)** Generalization index computed for context decoding across block-halves. Plots show mean ± s.e.m. block-half generalization index computed over bootstrap iterations for HPC (red), dACC (blue), and pre-SMA (green). **(D-F)** Same as **(A-C)**, but for context decoders trained and tested during the baseline period. **(G-J)** Cross-temporal generalization analysis control with ANOVA F-Statistic distribution matching between regions to ensure that increased temporal stability is not simply a consequence of stronger univariate context tuning at the single-unit level. eCDF of single-unit ANOVA F-statistics for each area are shown during the stimulus **(G)** and baseline **(I)** periods after performing distribution matching. Note: eCDFs for different regions are not clearly visible on the plots since they are practically identical after distribution matching. Cross-temporal generalization indices for the stimulus **(H)** and baseline **(J)** are recomputed using the matched distributions and presented as mean ± s.e.m. over bootstrap iterations for HPC (red), dACC (blue), and pre-SMA (green). **(K-L)** Cross-temporal generalization index for context computed on the control task variant where the trial response was given after a fixed delay instead of in a speeded manner. Plots show mean ± s.e.m. cross-temporal generalization index computed over bootstrap iterations for HPC (red), dACC (blue), and pre-SMA (green). Analysis is shown for both the stimulus period **(K)** and the baseline period **(L)**. Note: all p-values reported in this figure are computed by permutation test.

**Figure S5.**
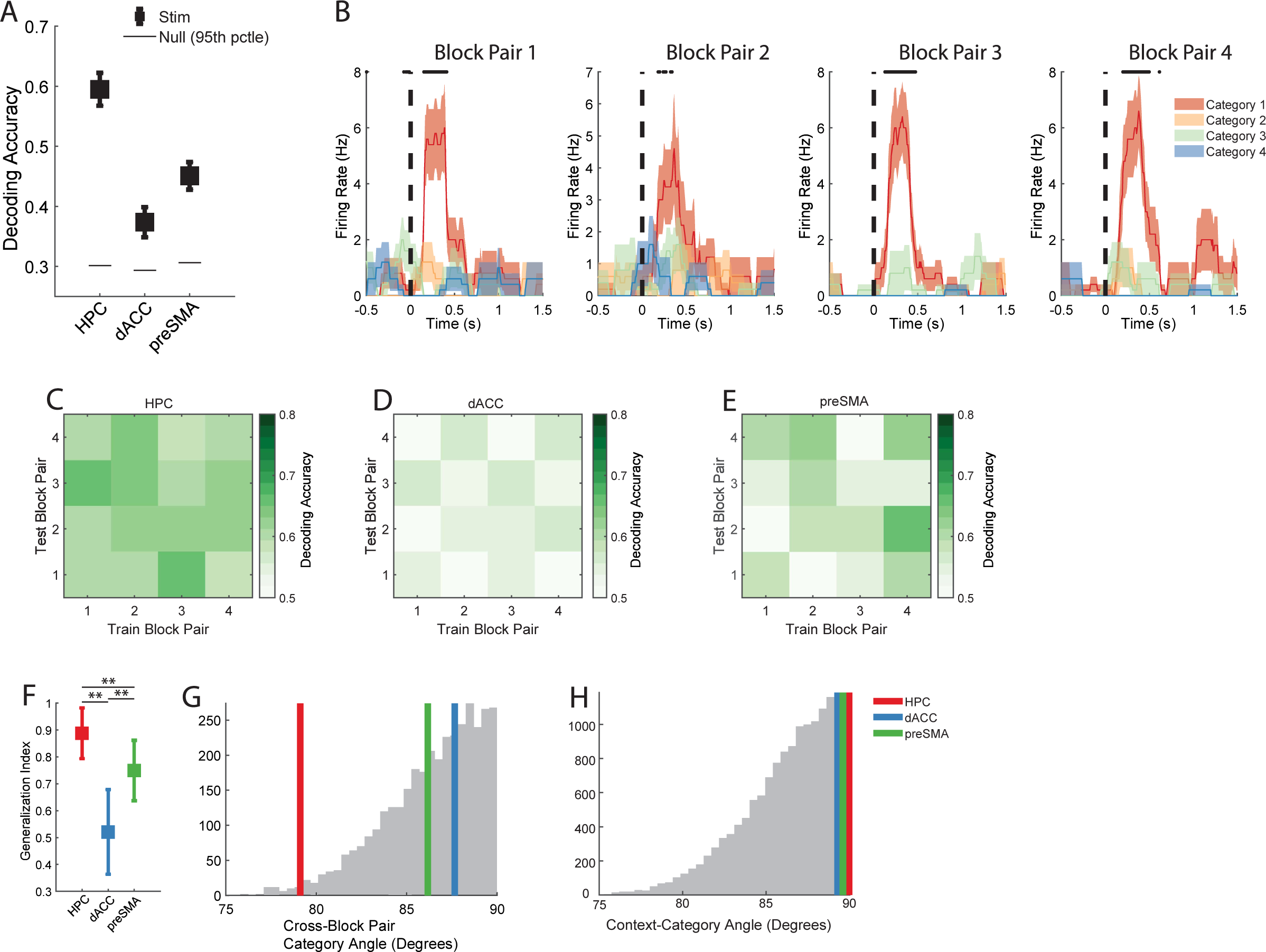
Hippocampal stimulus representation generalizes across time. **(A)** Decoding accuracy for semantic category of the presented stimulus during the stimulus period (0.2s to 1.2s following stimulus onset). Chance is 25% (4 categories). Plot shows mean decoding accuracy ± s.e.m. computed over bootstrap iterations. Horizontal black lines indicate 95^th^ percentile of shuffle null. **(B)** Example PSTHs for a single neuron exhibiting stable category selectivity (Category 1 preferred) plotted separately for every block pair in the experiment. All plotting conventions identical to those used for PSTHs in Fig. S1. **(C-E)** Cross-temporal decoding plots indicating decoding accuracy for decoders trained to decode image category from correct trials in adjacent block pairs during the stimulus period. All plotting conventions are identical to those in Fig. 2D. Cross-temporal decoding of image category is reported for HPC **(C)**, dACC **(D)**, and preSMA **(E)**. **(F)** Cross-temporal generalization index for image category computed using decoding accuracies reported in **(C-E)**. Plot shows mean decoding accuracy ± s.e.m. computed over bootstrap iterations for HPC (red), dACC (blue), and preSMA (red). P-values are computed by permutation test. **(G)** Angles computed between vectors normal to the hyperplanes of image category decoders for different block pairs. All plotting conventions are identical to those used in Fig. 2C. **(H)** Angles computed between vectors normal to the hyperplanes of image category decoders and the stimulus period context decoders for each region. All plotting conventions are identical to those used in Fig. 2C.

**Figure S6.**
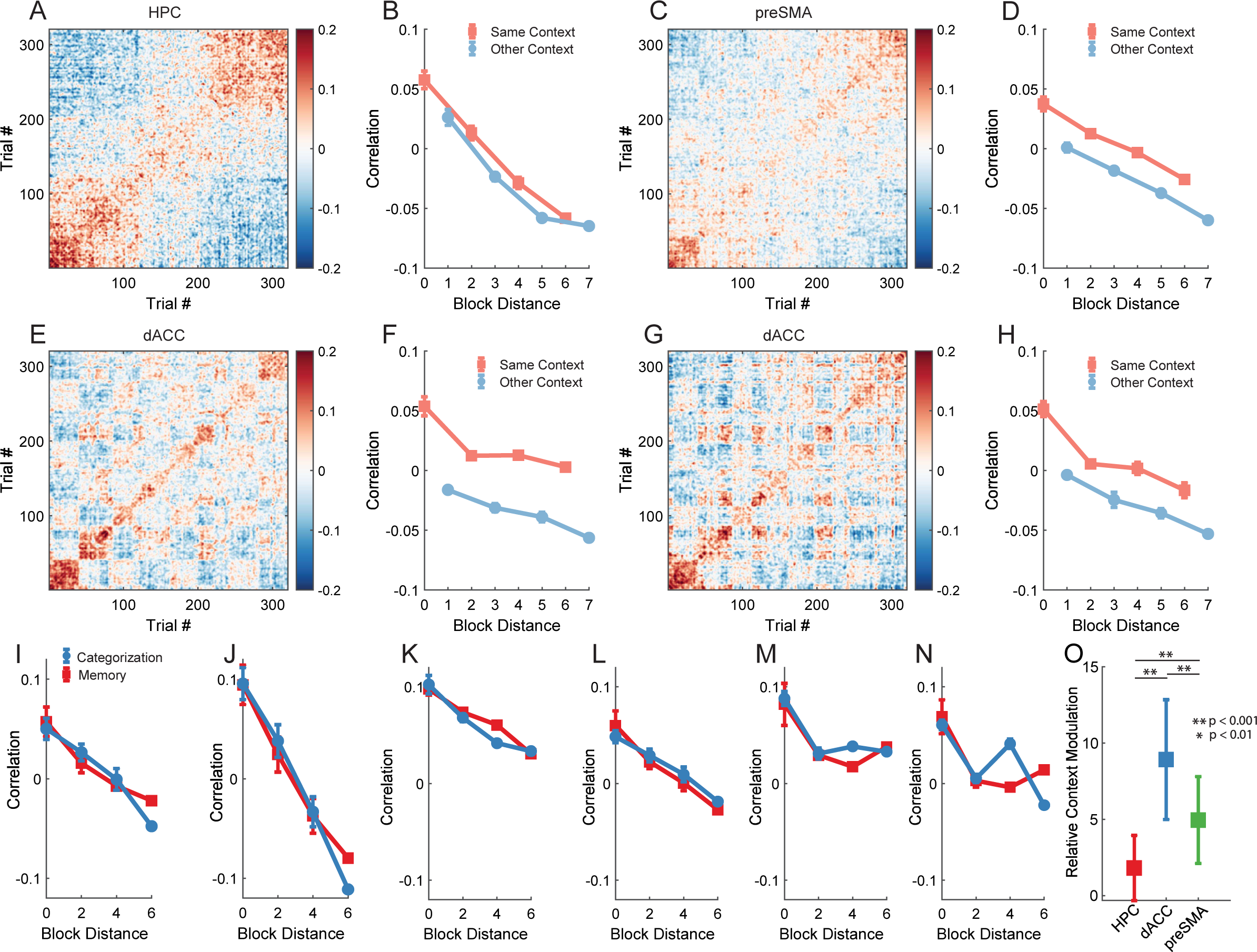
Additional population-vector autocorrelation analyses. **(A-H)** Trial-wise population vector autocorrelation plots and cross-block correlation curves shown for HPC during baseline **(A,B)**, preSMA during baseline **(C,D)**, and for the dACC during both stimulus **(E,F)** and baseline **(G,H)** periods. All plotting conventions identical to those used in Fig. 3. **(I-N)** Cross-block correlation curves reported separately for the categorization task (red) and the memory task (blue). Note in this case, since tasks always alternate, only even block distances can be computed and reported since there is no task block that is an odd number of blocks away from a block of the same task. All other plotting conventions identical to those used in Fig. 3B,D. Task-specific cross-block decorrelation curves are shown for the stimulus and baseline periods respectively in HPC **(I, J)**, preSMA **(K, L)**, and dACC **(M, N)**. **(O)** Relative context modulation reported for the three areas during the baseline period. Values are reported as mean ± s.e.m. over iterations of decorrelation curve estimation. P-values are computed by permutation test.

**Figure S7.**
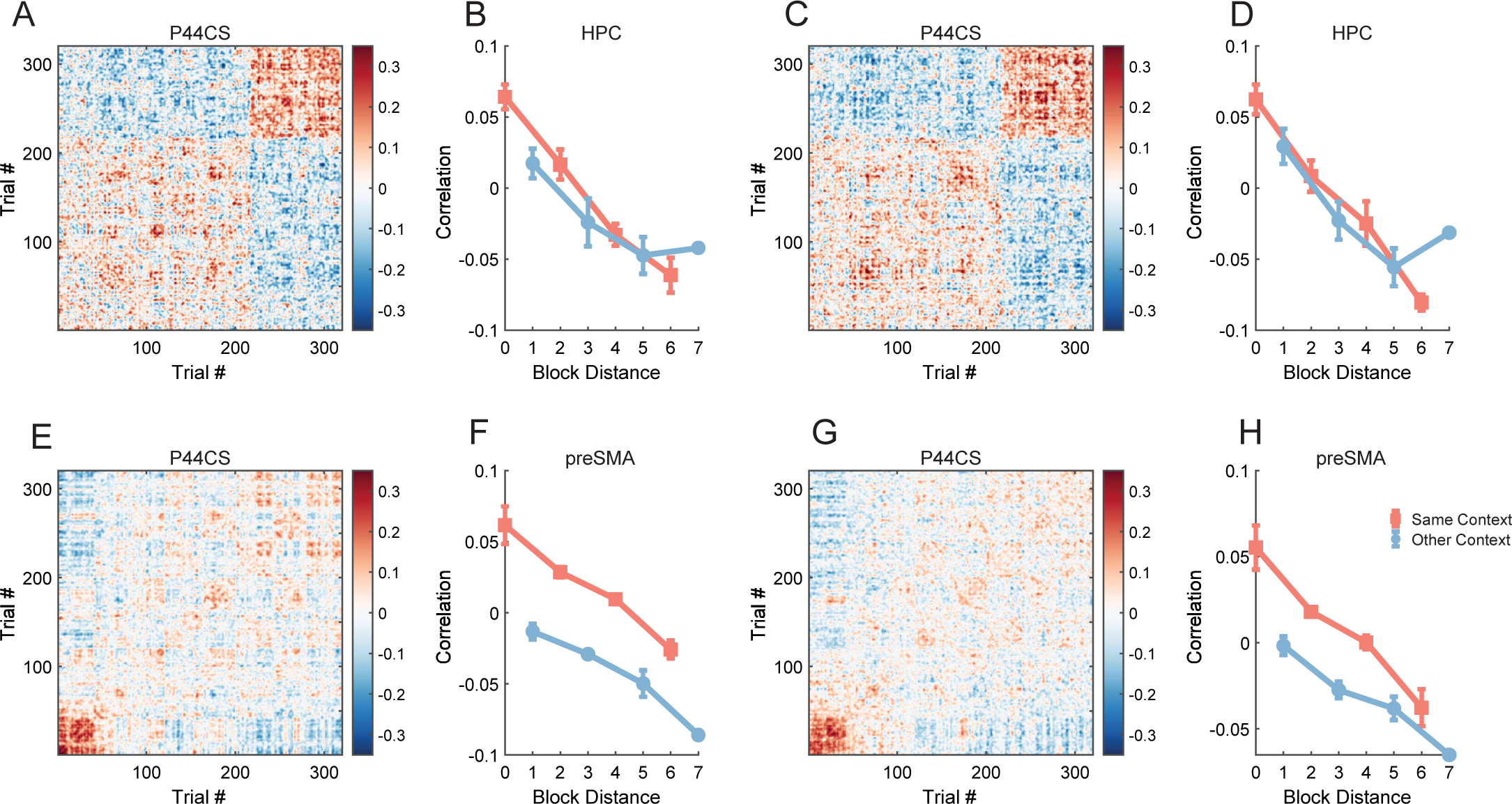
Single-subject recapitulation of temporal decorrelation effect. Recapitulation of area-dependent temporal decorrelation effect in neurons recorded in a single subject (P44CS). Trial-wise population vector autocorrelation plots and cross-block correlation curves are shown for HPC during stimulus **(A,B)** and baseline **(C,D)** periods and for preSMA during stimulus **(E,F)** and baseline **(G,H)** periods. All plotting conventions identical to those used in Fig. 3.

**Figure S8.**
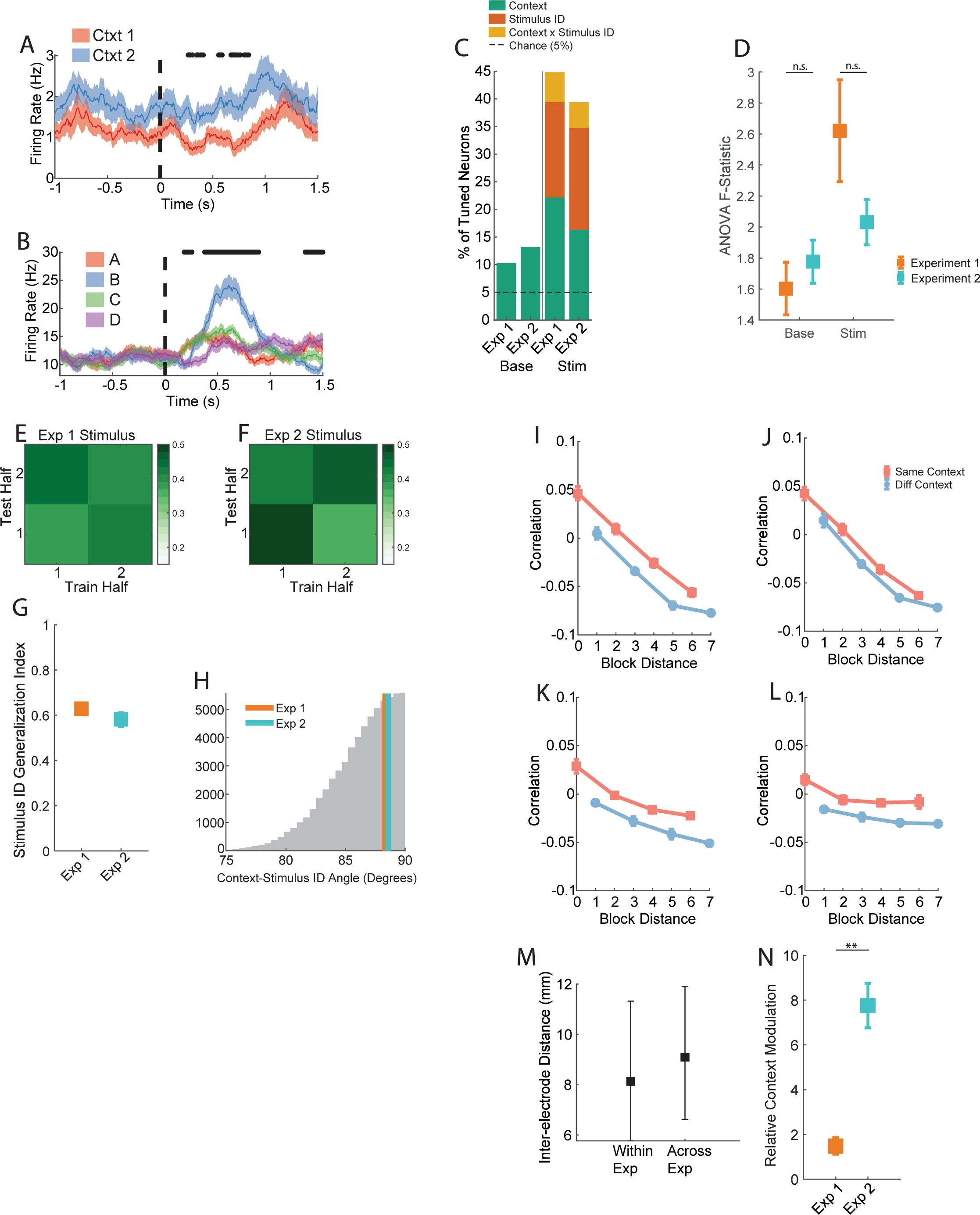
Single-unit properties and cross-temporal context decoding for Experiment 2. **(A-B)** Example PSTHs of two hippocampal neurons recorded from patients performing experiment 2. Neurons were modulated by the latent context variable **(A)** and by the identity of the stimulus presented on the screen **(B)**. Plotting conventions identical to those used in Fig. S1. **(C)** Percentage of hippocampal neurons that exhibit tuning to task variables during the Baseline (base, -1 to 0s prior to stimulus onset) and Stimulus periods (stim, 0.2 to 1.2 following stimulus onset). “Context” and “Stimulus” correspond to the task context variable and stimulus variable as applicable to each of the two experiments. 2-way (context x stimulus) ANOVAs are performed on firing rates for single neurons, where context and stimulus correspond to 2-level and 4-level categorical regressors respectively in both experiments. All other plotting conventions identical to those used in Fig. 1F. **(D)** ANOVA F-statistics for task context main effects shown for single neurons recorded in experiment 1 (left) and experiment 2 (right) during the baseline (red) and stimulus (blue) periods. Reported values are mean F-Statistic ± s.e.m. computed over neurons. P-values are computed by permutation test, and n.s. indicates p > 0.05. Cross-temporal decoding plots for image category in experiment 1 **(E)** and stimulus identity in experiment 2 **(F)** across experimental halves are shown for during the stimulus period. **(G)** Cross-temporal generalization index for the image category decoders reported for Experiment 1 (left) and stimulus identity decoders reported for Experiment 2 (right). Values reported are mean ± s.e.m. generalization index computed for the baseline (red) and stimulus (blue) periods of each task. P-values are computed by permutation test. **(H)** Angles computed between vectors normal to the hyperplanes of the image category decoder and the stimulus period context decoder in experiment 1 (orange), and of the stimulus ID decoder and the stimulus period context decoder in experiment 2 (teal). All plotting conventions are identical to those used in Fig. 2C. Cross-block correlation curves computed during the baseline **(I)** and stimulus **(J)** periods for experiment 1. Plots here are computed using the same data as those shown in Fig. 3B and **S5A**. **(K,L)** Same as **(I,J)** but for Experiment 2. **(M)** Distribution of all pair-wise inter-electro de distances within hemisphere computed within each experiment and pooled across the two experiment (Within Exp) and computed between all electrode pairs across the two experiments (Across Exp). Distances are reported as median with lower and upper error bars indicating 10^th^ and 90^th^ percentile respectively. **(N)** Relative context modulation reported for the hippocampus during the baseline period of experiments 1 and 2. Values are reported as mean ± s.e.m. over iterations of decorrelation curve estimation. P-value is computed by permutation test. **\*\*** indicates p < 0.001.

**Table S1. Tabulation of Patients, Behavior, and Neurons.**

Summary of patient information, the number of sessions performed for each experiment, the behavioral classification at the session level for experiment 2, and the number of recorded neurons per region per session. Patient behavior in experiment 2 is defined with respect to instances of high-level verbal instructions, where: Pre – “pre-instruction inference achieved”, NE – “Inference not exhibited”, post – “post-instruction inference achieved”, and N/A – “did not qualify for analysis”. Session behavior is defined with respect to performance on the first available inference trial, where: IA – “inference absent”, IP – “inference present”, X – “at or below chance non-inference performance”. Such definitions of patient behavior do not apply to experiment 1, and are listed as “N/A”.

